# The pangenome of the Arctic picoeukaryote *Micromonas polaris* resolves cryptic ecotypes and identifies a polar-specific gene repertoire

**DOI:** 10.64898/2026.11.16.765329

**Authors:** Victoria L. N. Jackson, Keith Harrison, Evelyne Derelle, Aurélie Chambouvet, Connie Lovejoy, Adam Monier

**Author notes:** Sorbonne Université/CNRS, UMR7144 Adaptation et Diversité en Milieu Marin, Station Biologique de Roscoff, Place Georges Teissier, 29680 Roscoff, France.

## Abstract

*Micromonas polaris* is the dominant picoeukaryote of the Arctic Ocean, where it sustains primary production across habitats from sea ice to open water to the subsurface chlorophyll maximum. How a single species maintains this ecological breadth has remained unclear. Standard phylogenetic markers such as 18S rRNA are nearly invariant within *M. polaris* and conceal any internal genomic heterogeneity the species may possess. Here we present a pangenome of 14 *M. polaris* strains spanning the Arctic basin, integrated with *Tara* Oceans metagenomes, that resolves the species into two diverged ecotypes invisible to 18S rRNA but recovered consistently by *prp8*, genome-wide SNPs, ANI, and orthogroup-based phylogenies. The two ecotypes coexist sympatrically, with one lineage consistently enriched deeper in the water column across the basin. *M. polaris* carries 990 polar-specific hierarchical orthogroups absent from temperate *Micromonas*, organised around one-carbon metabolism, endocytosis, transmembrane transport, and photo-oxidative stress response. Several components show phylogenetic signatures of horizontal acquisition from co-occurring marine taxa. Atlantification is bringing temperate *Micromonas* into Arctic margins seasonally, but the polar night likely limits their year-round persistence. *M. polaris* and its polar-specific toolkit appear positioned to persist through the dark season, suggesting Arctic margins will undergo seasonal turnover rather than stable replacement.

## Introduction

The Arctic Ocean is among Earth’s most extreme marine environments. Phytoplankton there must endure months of darkness, near-freezing temperatures, and marked seasonal swings in ice cover, light, and nutrient supply.^1,2^ Picocyanobacteria (*Prochlorococcus*, *Synechococcus*), which dominate the smallest phytoplankton size classes throughout temperate and tropical oceans, are largely absent.^3,4^ Green picoalgae of the order Mamiellales fill this vacant niche, and a single species, *Micromonas polaris*, frequently dominates summer Arctic picophytoplankton communities across the basin.^1,5–9^ *M. polaris* contributes substantially to Arctic primary production and carbon export^10^ across multiple heterogeneous habitats, from oligotrophic surface layers to the subsurface chlorophyll maximum (SCM), and from open water to the underside of sea ice.^11–13^ How a single species sustains dominance across such varied conditions remains unclear, and resolving this question is urgent: Atlantification is already pushing temperate *Micromonas* into Arctic margins,^10^ and the resilience of this keystone species will help determine whether Arctic planktonic communities retain their current composition or shift toward the temperate assemblages now appearing in Atlantic-influenced waters.^14–16^

Cryptic ecotypes (i.e., genetically distinct populations adapted to different environmental niches but indistinguishable by standard taxonomic markers) are increasingly recognised as a feature of marine picoeukaryote ecology. *Bathycoccus prasinos* and *B. calidus* were resolved as distinct cryptic species through genome-scale data after years of treatment as a single morphospecies.^17,18^ *Ostreococcus* clades partition the open ocean by depth and light, with surface-adapted and deep-adapted lineages occupying distinct vertical niches.^19^ In each case, fine-scale population structure underlies a phenotypically uniform species concept, with ecotypes responding differently to environmental forcing despite being collapsed into a single signal by 18S rRNA-based surveys.^19–23^ Recent taxonomic work has separated polar and temperate *Micromonas* lineages, previously conflated under *M. pusilla*, into four distinct species (*M. polaris*, *M. commoda*, *M. pusilla*, and *M. bravo*),^24^ confirming that cryptic speciation is a hallmark of the genus. *M. polaris* is currently understood to be a homogeneous and narrowly cold-adapted species lacking the intraspecific diversity seen in its sister lineages.^1,25,26^ However, this consensus rests on conventional markers that have systematically failed to resolve fine-scale structure in related species.

Polar phytoplankton must function in conditions that pose distinct challenges relative to the temperate ocean: extended darkness during the polar night, cold temperatures that slow enzymatic kinetics and increase membrane rigidity, sharp seasonal transitions in irradiance that combine increased light with low temperatures, fluctuating salinities during ice freeze up and melt seasons,^27,28^ and oligotrophic conditions interrupted by intense bloom events.^29^ Genome sequencing of cultured polar microalgae has begun to reveal cold-adapted genomic features, including ice-binding proteins, expanded gene families for photoprotection and DNA repair, and condition-specific allele or gene expression under freezing, darkness, and variable irradiance.^30–33^ Comparative work has further highlighted horizontal gene transfer (HGT) from co-occurring marine taxa as a recurrent feature of Arctic algal genome evolution,^34^ with sea ice proposed as a particular hotspot for lateral gene exchange among algal communities.^35,36^ Whether *M. polaris* carries an analogous polar-specific gene complement and whether HGT from co-occurring marine taxa contributes to that complement remain untested and impossible to assess without genomic data. As yet, no published genome of *M. polaris* exists, in contrast to its temperate congeners *M. commoda* and *M. pusilla,* which have multiple sequenced genomes.^37,38^

Here we combine pangenomics with metagenomic recruitment to test two hypotheses about *M. polaris*. First, the species comprises genetically distinct ecotypes that partition the heterogeneous Arctic habitats it occupies, with the cryptic population structure invisible to 18S rRNA but resolvable with whole-genome data and finer-scale phylogenetic markers. Second, *M. polaris* possesses a polar-specific gene repertoire distinguishing it from temperate *Micromonas* species. To test these hypotheses, we sequenced and assembled the genomes of 14 *M. polaris* strains spanning four Arctic regions and three habitat categories, constructed a *Micromonas* pangenome integrating these strains with the existing temperate *Micromonas* genomes, and recruited the resulting clade-level genomic data against *Tara* Oceans metagenomes from the Arctic basin to test ecotype distribution at the population level.

## Results

### A pan-Arctic strain panel for *M. polaris* pangenomics

To capture as much of the within-species variation in *M. polaris* as the available cultured diversity allows, we assembled a panel of 14 strains isolated from four Arctic regions over four oceanographic campaigns conducted between 1998 and 2016 (Fig. 1a; Extended Data Table 1). The panel includes the *M. polaris* type strain RCC807, isolated from the Pikialasorsuaq (formerly the North Water Polynya in Northern Baffin Bay)^5^; seven strains from the MALINA expedition in the Beaufort and Chukchi Seas^39^; two strains from the MicroPolar project north of Svalbard^40,41^; and four strains from the GreenEdge Ice Camp at Qikiqtarjuaq, Baffin Bay.^42,43^ Strains were isolated across a wide range of seasons (March to August), depths (0 to 500 m), and sea ice conditions, providing a broad sampling of the *M. polaris* niche.

**Figure 1.**
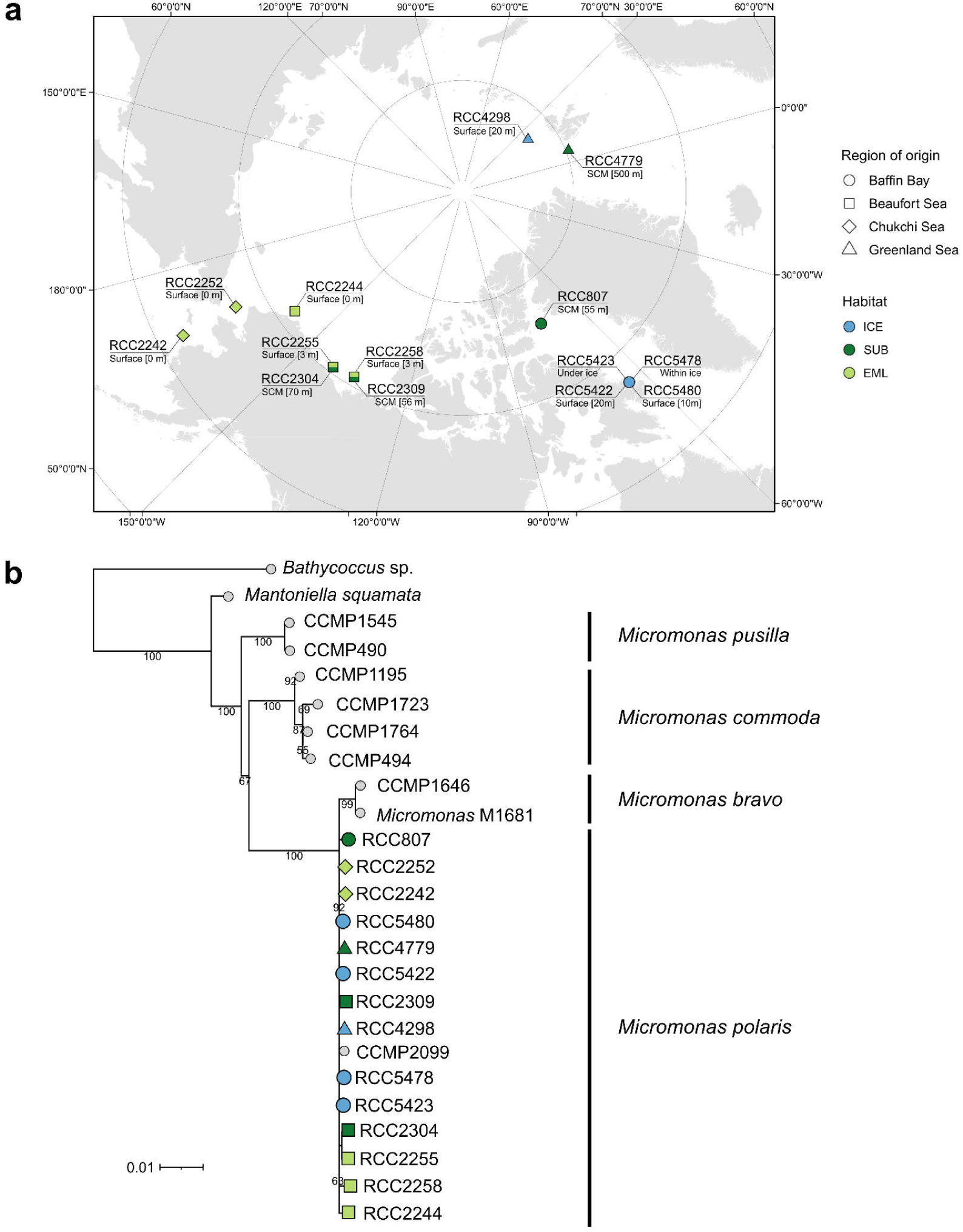
Geographic origin and 18S rRNA phylogeny of the 14 *Micromonas polaris* strains analysed in this study. **a)** Arctic map showing the isolation location, habitat, and isolation depth of each strain. Symbol shape denotes the Arctic region of origin (circle, Baffin Bay; square, Beaufort Sea; diamond, Chukchi Sea; triangle, Greenland Sea); symbol fill denotes the habitat category (light blue, sea ice; light green, euphotic mixed layer; dark green, subsurface). Strain names and isolation depths are annotated for each site. **b)** Maximum-likelihood phylogenetic tree of *Micromonas* spp. and the related Mamiellales *Bathycoccus* sp. and *Mantoniella squamata*, inferred from a 1,659-site alignment of near-full-length 18S rRNA gene sequences using the GTR+F+G4 model. The 14 *M. polaris* strains analysed here are shown with the same coloured symbols as in panel a; previously published sequences are shown as small grey circles. Bootstrap support values from 1,000 non-parametric replicates are shown for internal nodes with support ≥50. Scale bar represents substitutions per site. Four *Micromonas* species are labelled: *M. pusilla*, *M. commoda*, *M. bravo*, and *M. polaris*.

Standard isolation labels in culture collections (e.g., “surface” or “under ice”) capture isolation depth and physical context but do not consistently reflect the ecological setting in which a strain was sampled. A strain isolated at 20 m through landfast sea ice in mid-summer Baffin Bay (RCC5422) and a strain isolated at 0 m in open water in August Beaufort Sea (RCC2258) share the “surface” or “shallow” label but represent fundamentally different habitats. To preserve this distinction in our analyses, we initially classified each strain into one of three habitat categories defined by environmental context (Fig. 1a; Extended Data Table 1).

#### Sea ice-influenced

Strains isolated within sea ice, immediately under sea ice, or in ice-free water within heavily ice-covered regions during winter or pre-bloom conditions were designated sea ice-influenced. This category captured strains associated with sea ice cover encompassing samples from within and beneath ice, or in open water adjacent to sea ice. RCC5478 was isolated from melted ice cores; RCC5423, RCC5480, and RCC5422 were isolated from the water column directly beneath landfast sea ice at the GreenEdge Ice Camp, and RCC4298 was isolated in early March from waters of the marginal ice zone north of Svalbard in the Greenland Sea, where 2014 ice cover was unusually heavy.^40,41^

#### Euphotic mixed layer

Strains isolated from the upper open-water column during summer in conditions consistent with an established seasonal mixed layer within the euphotic zone. The five strains in this category (RCC2242, RCC2244, RCC2252, RCC2255, RCC2258) were all isolated from 0 to 3 m in the Beaufort or Chukchi Seas in July or August 2009, well after ice retreat and when light levels would be greatest.

#### Subsurface

Strains isolated from below the seasonal mixed layer in open water were considered subsurface. This category included three strains isolated from the subsurface chlorophyll maximum (SCM) in the Beaufort Sea or Baffin Bay (RCC2304, RCC2309, RCC807, isolation depths 55 to 70 m) and one strain isolated at 500 m in the Greenland Sea (RCC4779).

The five strains in the sea ice-influenced category, the five in the euphotic mixed layer, and the four subsurface strains together spanned the major water-column habitats *M. polaris* is known to occupy in the Arctic Ocean. We use these categories throughout the study to interpret strain-level findings and to map them onto the population-scale patterns recovered from environmental metagenomes.

A first phylogenetic placement of the 14 strains using near-full-length 18S rRNA gene sequences (∼1,680-1,690 bp amplicons; 1,659-site alignment after trimming; Extended Data Table 2) recovered the *M. polaris* strains as a tightly clustered group within the Mamiellales (Fig. 1b), with effectively no resolvable diversity (0.1% sequence divergence) within *M. polaris* despite the geographic, seasonal, and habitat range represented. This near-invariance was consistent with previous reports that 18S rRNA cannot resolve fine phylogenetic structure within this marine picoeukaryote species^1,5,26^ and motivated us to investigate phenotypic and genomic variation as alternative routes into the diversity of *M. polaris*.

### Contrasting phenotypes and cryptic diversity

While maintaining cultures of the panel under standard low-light conditions, we noted that strains reached high cell density at different rates. To test whether these growth differences tracked habitat of origin as designated above, we monitored growth and photosynthetic quantum yield (*F*_v_/*F*_m_) of one representative strain per habitat for 14 days under high light (HL; 60 µmol photons m⁻² s⁻¹) and low light (LL; 1.5 µmol photons m⁻² s⁻¹) at 4 °C. RCC2258 represented the euphotic mixed layer category, RCC5478 the sea ice-influenced (within-ice isolate) category, and RCC2309 the subsurface category (SCM, 56 m).

*M. polaris* strains differed in growth rates and photophysiology under the two light conditions. The euphotic mixed layer strain (RCC2258) and the subsurface strain (RCC2309) both grew faster and maintained higher *F*_v_/*F*_m_ under LL than HL, with RCC2309 outpacing RCC2258 in LL (Fig. 2). The sea ice strain (RCC5478) tolerated both light regimes, although its photophysiology and growth still favoured LL, consistent with the variable irradiances experienced beneath sea ice.^44^ A Kruskal-Wallis test detected an effect of light treatment on growth rate (H(1)=4.08, *p*=0.04), and the only significant pairwise difference between strains separated RCC2258 and RCC5478 (Wilcoxon, *p*-adj=0.007). With only one strain per habitat, these observations are only suggestive, but show that strains from different habitats differ in photophysiology when grown under identical laboratory conditions despite the high similarity of their 18S rRNA gene.

**Figure 2.**
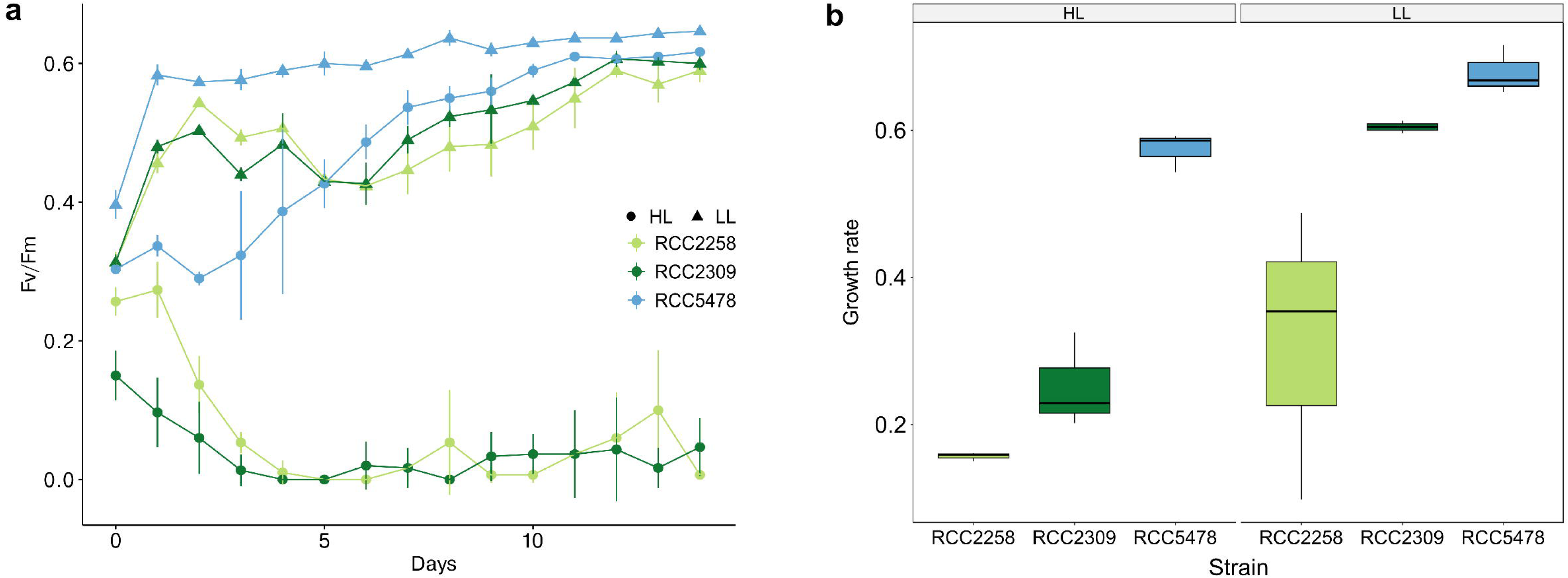
Photophysiology of three *M. polaris* strains representing distinct Arctic habitat categories under contrasting light regimes. Three strains were selected to represent the three habitat categories from which the strain panel was isolated: RCC2258 (euphotic mixed layer; Beaufort Sea, 3 m), RCC2309 (subsurface; Beaufort Sea, 56 m, SCM), and RCC5478 (sea ice-influenced; Baffin Bay, within-ice). Strains were grown at 4°C under high light (HL; 60 μmol photons m⁻² s⁻¹) and low light (LL; 1.5 μmol photons m⁻² s⁻¹) in n=3 biological replicates per strain and light combination. **a)** Quantum yield of photosystem II (*F*_v_/*F*_m_) measured daily over a 14-day time course. Symbols denote light treatment (filled circles, HL; filled triangles, LL); colours denote strain (light green, RCC2258; dark green, RCC2309; blue, RCC5478). Error bars represent standard deviation across biological replicates. **b)** Growth rates calculated from cell counts on days 6–13 of the growth curve, with each strain and light combination represented by a boxplot.

However, the habitat categories of the three strains were not congruent with what may have been predicted from the light conditions during the original isolation of the strains (Fig. 2). To search for the genetic basis of this phenotypic variation, we turned to *prp8*, a less conserved phylogenetic marker that has resolved cryptic diversity in the related Mamiellales picoeukaryote *Bathycoccus*.^45^ We designed *M. polaris*-specific primers targeting a 975 bp region in a conserved exon of the gene (Extended Data Fig. 1; Extended Data Table 2) and amplified and sequenced the corresponding fragment from all 14 strains. The *prp8* phylogeny resolved *M. polaris* into two intraspecific subclades, separated by 6 bp in a 911 bp alignment of their consensus sequences (Fig. 3). Eight strains formed one subclade and five formed the other; RCC4298 branched separately with weak support (<50%).

**Figure 3.**
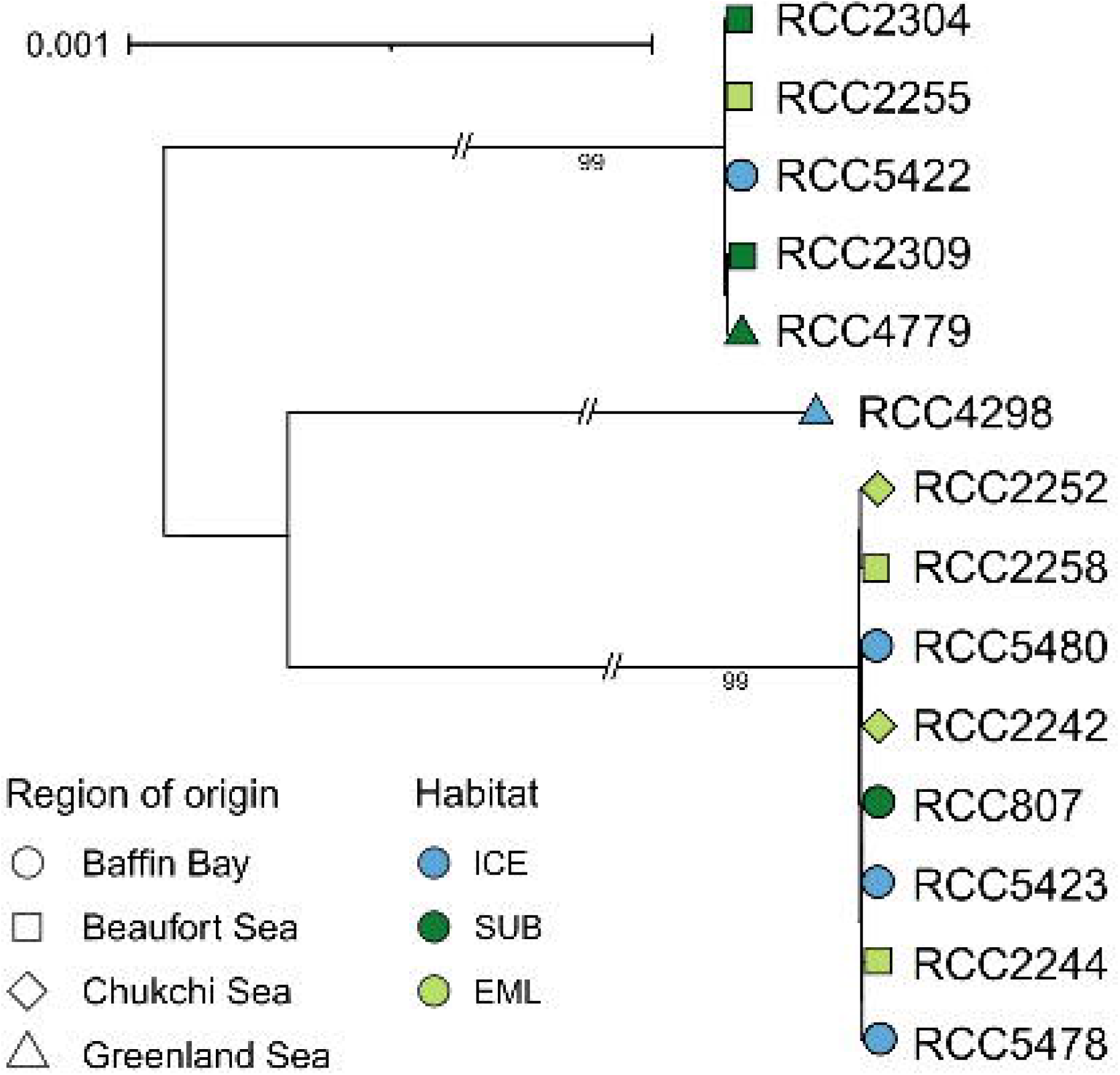
The *prp8* phylogeny resolves *M. polaris* into two intraspecific subclades. Maximum-likelihood midpoint-rooted phylogenetic tree of the 14 *M. polaris* strains analysed in this study, inferred from a 903-site alignment of *prp8* gene fragments under the model F81+F. The amplified *prp8* region (975 bp) was trimmed to remove poorly aligned positions, yielding a 903-site alignment for tree inference. Strain symbol shape denotes Arctic region of origin (circle, Baffin Bay; square, Beaufort Sea; diamond, Chukchi Sea; triangle, Greenland Sea); symbol fill denotes habitat type (light blue, sea ice-influenced; dark green, subsurface; light green, euphotic mixed layer). Two intraspecific clades are resolved with strong support (eight strains in one clade and five in the other; bootstrap 99 in both cases). RCC4298 (Greenland Sea, surface, 20 m) branches separately with bootstrap support below 50 and is placed as a divergent member of the second clade. Bootstrap support values from 1,000 non-parametric replicates are shown for internal nodes with support ≥50. Branch break symbols (//) indicate that long internal branches between clades are shown compressed; the relative within-subclade branch lengths are drawn to scale. Scale bar represents 0.001 substitutions per site.

Strain subclades based on the *prp8* phylogeny were not completely congruent with the habitat of origin categories (Fig. 4). One subclade contained four euphotic mixed layer strains (RCC2242, RCC2244, RCC2252, RCC2258), three sea ice-influenced strains (RCC5423, RCC5478, RCC5480), and the type strain RCC807 (subsurface). The other subclade contained three subsurface strains (RCC2304, RCC2309, RCC4779), one sea ice-influenced strain (RCC5422), and one euphotic mixed layer strain (RCC2255), with RCC4298 (sea ice-influenced) branching separately. Strains from the same Arctic region were distributed across both subclades wherever sampling allowed the comparison (e.g. Baffin Bay strains appeared in both subclades), indicating that the separation was not driven by geography. The *prp8* phylogenetic structure, invisible using 18S rRNA, together with the phenotypic diversity among strains from the HL/LL experiment, motivated a genome-scale investigation to determine whether the subclades reflected deeper genomic differentiation with ecologically relevant consequences for *M. polaris*.

**Figure 4.**
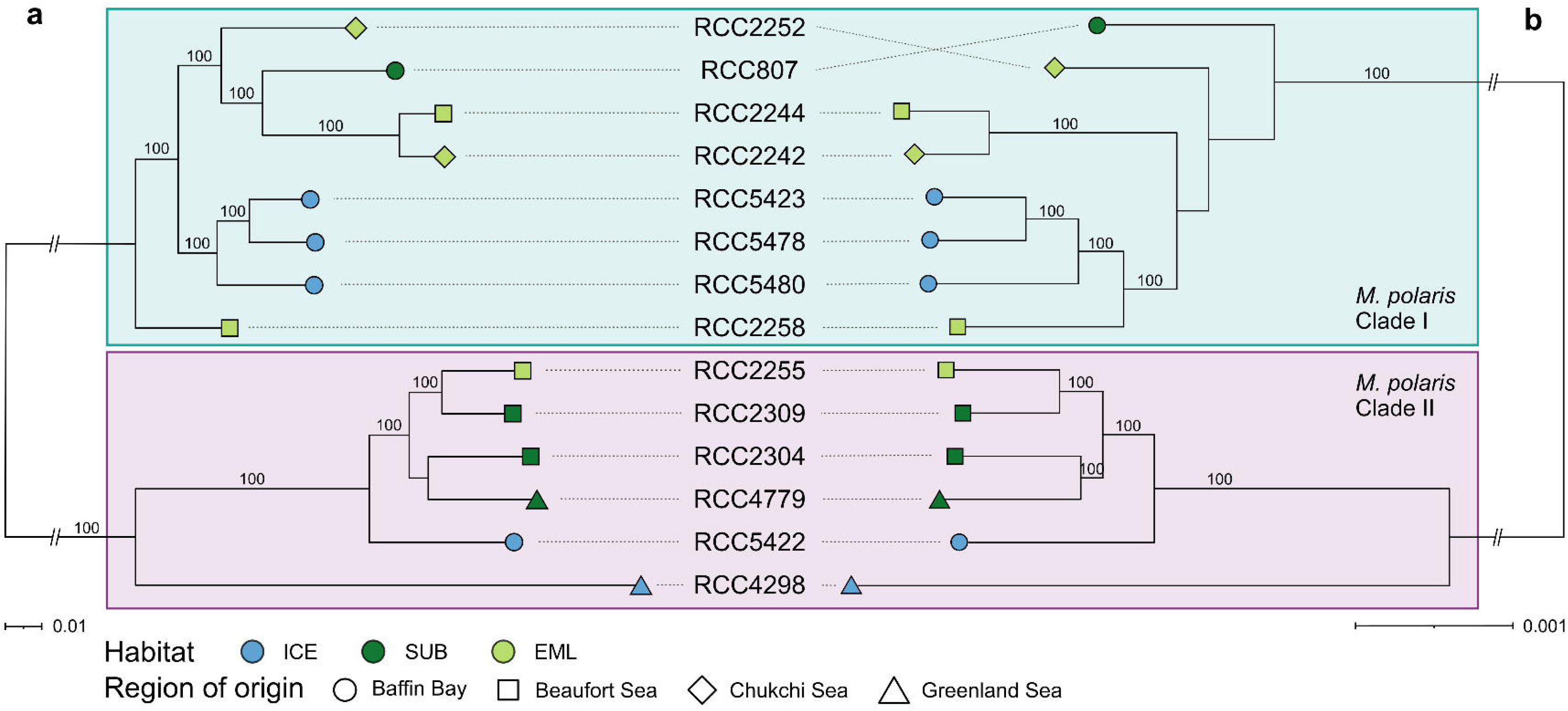
Genome-wide SNP and orthogroup phylogenies recover the same two-clade structure within *M. polaris*. Two independent phylogenetic reconstructions of the 14 *M. polaris* strains, both recovering Mpol Clade I (eight strains, teal background) and Mpol Clade II (six strains, pink background). Strains symbol shape denotes Arctic region of origin (circle, Baffin Bay; square, Beaufort Sea; diamond, Chukchi Sea; triangle, Greenland Sea); symbol fill denotes habitat type (light blue, sea ice-influenced; dark green, subsurface; light green, euphotic mixed layer). **a)** Maximum-likelihood phylogenetic tree inferred from a 329,175-site SNP matrix generated by mapping all 14 strains against the RCC807 reference assembly and calling single-nucleotide variants. Bootstrap support values from 1,000 non-parametric replicates are shown only at internal nodes with maximum support (100). **b)** Species tree generated by OrthoFinder v3 (inferred with STAG^131^) using a concatenated sequence alignment of 3772 single-copy orthogroups present in all *Micromonas* spp. genomes used. The two analyses recover the same Clade I / Clade II topology. RCC4298, which branched separately on the *prp8* tree (Fig. 3), is placed within Clade II as a divergent member in both genome-wide analyses, on a long branch reflecting its substantial sequence divergence from other Clade II strains. Scale bars represent substitutions per site (a, 0.01; b, 0.001); the order-of-magnitude difference reflects the higher mutation rate captured at the SNP level relative to the orthologous protein-coding regions used by OrthoFinder.

### Genome-level analyses

#### Two *M. polaris* ecotypes recovered consistently across genome-wide analyses

The nuclear genomes of all 14 strains were sequenced and assembled using a hybrid short- and long-read sequencing approach (Illumina and Oxford Nanopore platforms). Final assemblies were highly contiguous and complete; each comprised 19 chromosomes, ranged from 22.69 to 23.09 Mb, had GC content near 59%, and recovered 97.5 to 98.2% of universal single-copy orthologs (BUSCO, Table 1). Predicted gene counts ranged from 10,693 (RCC807) to 11,121 (RCC4298). Pulsed-field gel electrophoresis of RCC807 prior to assembly was consistent with the 19-chromosome architecture (Extended Data Fig. 2).

Genome-wide phylogenomics confirmed the *prp8* split and revealed two deeply diverged lineages within *M. polaris*. A SNP-based maximum-likelihood phylogeny built from 329,175 sites mapped to RCC807, an OrthoFinder species tree built from shared orthogroups, and PCA of population structure all recovered the same two-way division (Fig. 4; Extended Data Fig. 3). Average nucleotide identity (ANI) was higher within than between the two clades (within Clade I: 99.66 - 99.92%, mean = 99.72%; within Clade II: 99.28 - 99.84%, mean = 99.63%; between clades: 98.91 - 99.07%; mean = 99.01%; Extended Data Fig. 4), consistent with the phylogenetic split. Strain assignments matched the *prp8* result exactly, validating *prp8* as a clade marker for *M. polaris* and confirming that the division reflected substantial genome-wide differentiation rather than localised divergence at a single locus. We hereafter refer to these lineages as Mpol Clade I and Mpol Clade II.

The two clades partition the *M. polaris* niche by a less nuanced habitat category rather than by geography. Mpol Clade I (n=8) was concentrated in shallow euphotic and ice-influenced habitats, with four euphotic mixed layer strains (RCC2242, RCC2244, RCC2252, RCC2258) and three sea ice-influenced strains (RCC5423, RCC5478, RCC5480). The exception was the type strain RCC807, which was considered subsurface. In contrast, half of the 6 Mpol Clade II strains originated from subsurface habitats spanning the subsurface chlorophyll maximum and below (RCC2304, RCC2309, RCC4779, with isolation depths from 56 to 500 m). The clade also included two sea ice-influenced strains (RCC4298, RCC5422) and one euphotic mixed layer strain (RCC2255). Both clades co-occurred wherever sampling allowed the comparison: Baffin Bay and the Beaufort Sea each yielded strains from both clades, indicating sympatry rather than allopatric divergence. The apparent absence of one clade or the other from the Chukchi and Greenland Seas more likely reflects the small number of strains from each region (n=2) than genuine biogeographic structure. We hereafter treat Mpol Clade I and Mpol Clade II as candidate ecotypes, with Clade I associated with the higher light in the euphotic mixed layer and just under the ice cover of ice-influenced habitats, and Clade II associated with subsurface and from deeper water (>20 m depth) of ice-influenced habitats (Table S1) where light levels would be lower.

RCC4298 occupied a divergent position within Mpol Clade II across multiple analyses. It formed a long-branched, basal lineage of Mpol II in both the SNP and OrthoFinder phylogenies (Fig. 4), and shared the lowest within-clade ANI with other Clade II members (Extended Data Fig. 4). Despite this divergence, all genome-scale analyses placed it within Mpol Clade II rather than as a separate lineage. RCC807, the only Mpol Clade I strain isolated below the seasonal mixed layer, showed no comparable divergence; it sat well within the clade in both genome trees and showed no long-branch effect. Its anomalous habitat assignment may reflect the distinctive physical regime of the April 1998 North Water Polynya, where convective mixing and ice-edge dynamics blur the surface-versus-subsurface distinction, rather than a SCM-adapted phenotype.

#### Mobile and viral elements concentrate in outlier chromosomes

Mamiellales genomes typically contain two structurally distinct outlier chromosomes, a big outlier chromosome (BOC) carrying a low-GC, repeat-rich BOC1 region, and a small outlier chromosome (SOC), that differ in gene content, recombination, and drive changes in viral susceptibility not seen elsewhere in the nuclear genome.^46–48^ *M. polaris* genomes followed this pattern, but the size and chromosomal location of both outlier features varied considerably among strains (Fig. 5). BOC1 ranged from 271 kb (RCC2309) to 430 kb (RCC5478) and was located on chromosome 1 or 3 in most strains (n=10), with the remainder on chromosomes 2 (n=3) or 4 (n=1). SOCs sat on either chromosome 18 (n=10) or 19 (n=4).

**Figure 5.**
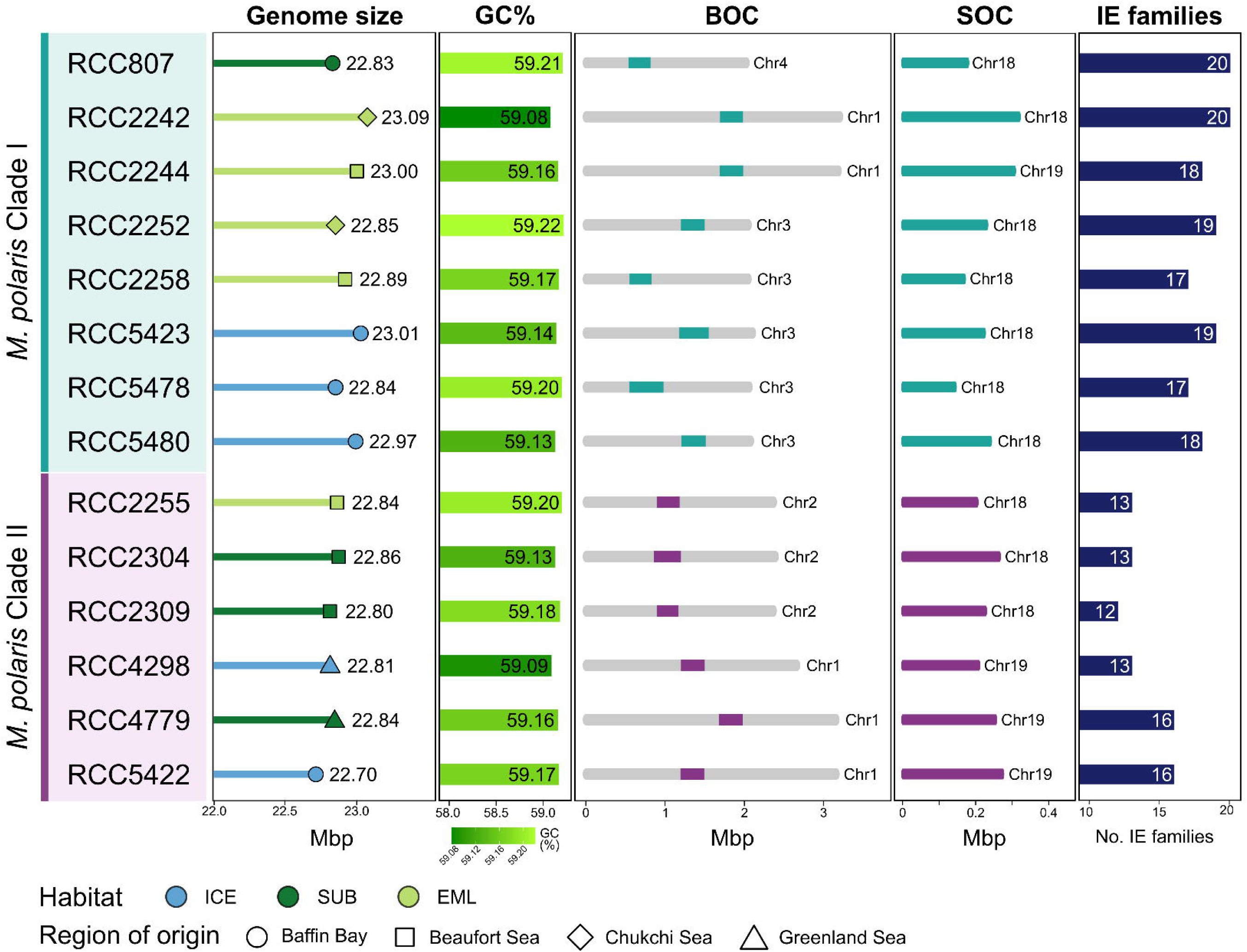
Genome architecture and mobile element content of the 14 *M. polaris* strains, grouped by clade. Mpol Clade I (top eight strains, teal background) and Mpol Clade II (bottom six strains, pink background) are defined by orthogroup-based phylogenetic analysis. Each row represents one strain, symbol shape denotes Arctic region of origin (circle, Baffin Bay; square, Beaufort Sea; diamond, Chukchi Sea; triangle, Greenland Sea); symbol fill denotes habitat category (light blue, sea ice-influenced; light green, euphotic mixed layer; dark green, subsurface). **Genome size:** total assembly size in Mbp. **GC%:** mean genome-wide GC content. **BOC:** schematic of the big outlier chromosome (BOC), a structurally distinct chromosome conserved across Mamiellales and characterised by elevated mobile-element content. The BOC is carried on different chromosomes across strains (Chr1 to Chr4, labelled at right of each schematic). The coloured region within each BOC marks the BOC1 region, the variable low-GC subregion that concentrates transposable, introner, and viral elements (teal, Clade I; magenta, Clade II). **SOC:** schematic of the small outlier chromosome (SOC), a second structurally distinct chromosome present in all *M. polaris* genomes (Chr18 or Chr19, labelled at right; teal, Clade I; magenta, Clade II). **IE families:** number of distinct introner element families identified per genome (range 17–20 in Clade I, 12–16 in Clade II).

Mobile and viral elements were distributed non-randomly across these compartments (Extended Data Fig. 5). Introner element (IE) families ranged from 12 to 20 per genome and were significantly depleted in both outlier chromosomes relative to standard genomic regions (BOC mean 0.002/kb, SOC mean 0.0003/kb, standard 0.13/kb; Wilcoxon, both *p*<0.0001). Transposable elements showed the opposite pattern, accumulating in SOCs (0.1/kb) relative to BOCs (0.04/kb; *p*<0.01) and standard regions (0.01/kb; *p*<0.0001). Viral elements were enriched in SOCs (0.03 hits/kb) compared with BOCs (0.003 hits/kb; *p*<0.01) and standard regions (0.01 hits/kb; *p*<0.0001). The outlier chromosomes thus represent distinct genomic compartments in *M. polaris*, with SOCs in particular accumulating mobile and viral DNA, a pattern consistent with their proposed roles as sites of accelerated evolution and host-virus interaction in other Mamiellales.^47,49^

#### IE distribution patterns track *M. polaris* diversity

The number of IE families in Mpol Clade I ranged from 17 to 20 and in Mpol Clade II from 12 to 16 (Fig. 5). Clustering and dereplication of all the IE families identified across the *M. polaris* genomes yielded a total of 66 IE clusters, of which 21 were present in only one strain (Extended Data Fig. 6). Hierarchical clustering of the *M. polaris* genomes based on the abundance profiles of IE clusters in each *M. polaris* genome also recovered the same two-clade structure as the other genome-level analyses.

#### The *M. polaris* polar-specific core is over twice the size of the temperate *Micromonas*-specific core

To identify genomic features distinguishing *M. polaris* from its temperate relatives and to test whether Mpol Clade I and Mpol Clade II differ in functional content, we constructed a *Micromonas* pangenome from the 14 *M. polaris* genomes alongside *M. pusilla* CCMP1545 and *M. commoda* RCC299. OrthoFinder grouped predicted proteins into 10,570 hierarchical orthogroups (HOGs), of which 6,762 were core to all *Micromonas* genomes (Fig. 6a, Table 2). The *M. polaris* pangenome alone comprised 10,129 HOGs of which 990 formed the polar-specific core (Fig. 6a, Table 2).

**Figure 6.**
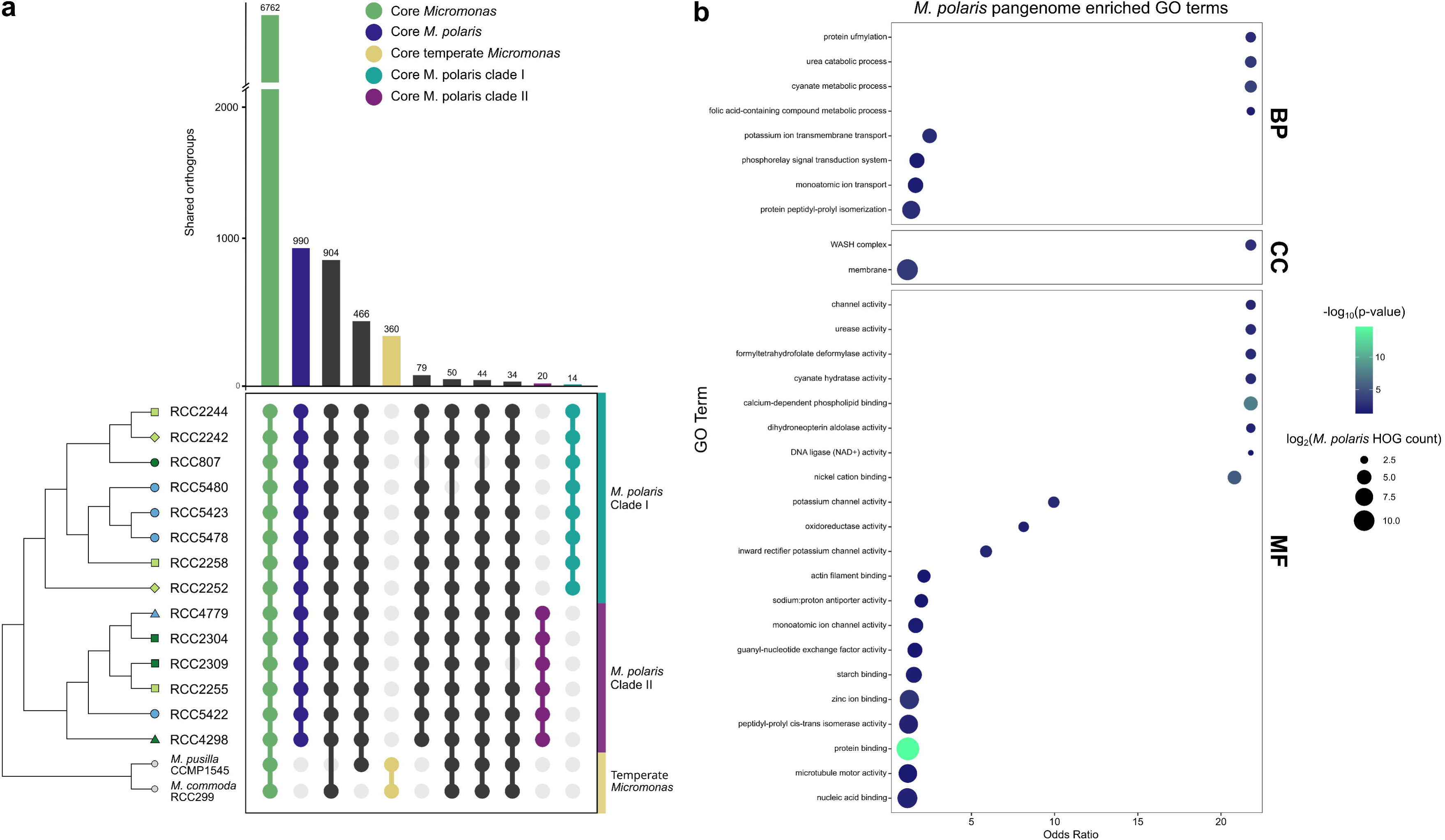
The *M. polaris* polar-specific gene complement is an order of magnitude larger than the temperate-specific complement. **a)** Upset plot of hierarchical orthogroup (HOG) intersections among the 14 *M. polaris* strains and the temperate species *M. pusilla* CCMP1545 and *M. commoda* RCC299, generated by OrthoFinder v3. Bars represent HOG counts at each intersection (vertical axis broken to accommodate the large core *Micromonas* set); the dot matrix below indicates which genomes contribute to each intersection. Coloured bars highlight key intersections: core *Micromonas* (6,762 HOGs shared by all 16 genomes; green), core *M. polaris* (990 HOGs shared by all 14 *M. polaris* genomes and absent from temperate *Micromonas*; dark blue), core temperate *Micromonas* (360 HOGs shared by *M. pusilla* and *M. commoda* and absent from *M. polaris*; yellow), core Mpol Clade I (14 HOGs shared by all eight Clade I strains and absent from all other genomes; teal), and core Mpol Clade II (20 HOGs shared by all six Clade II strains and absent from all other genomes; magenta). The polar-specific core (990 HOGs) is several-fold larger than the temperate-specific core (360 HOGs). Strain identifiers at left use the same coloured symbols as in Fig. 1; clade backgrounds are: teal, Clade I, and pink, Clade II. **b)** GO term enrichment of the *M. polaris* pangenome compared to the temperate *Micromonas* pangenome, shown for biological process (BP), cellular component (CC), and molecular function (MF) categories in separate panels. Each bubble represents one significantly enriched GO term, with bubble size scaled to log₂ of the *M. polaris* HOG count assigned to that term, bubble colour scaled to -log₁₀ of the enrichment p-value (colour scale at right), and horizontal position showing the odds ratio of the enrichment test. Enriched terms span GO categories relating to one-carbon and folate metabolism, endosomal trafficking, transmembrane ion transport, and protein turnover.

**Table 2.** *Micromonas* hierarchical orthogroup (HOG) repertoires. HOG counts from the pangenome of the 14 *M. polaris* strains and the temperate species *M. pusilla* CCMP1545 and *M. commoda* RCC299 (Fig. 6a). Pangenome totals give the number of HOGs present in at least one genome of the indicated set. Core sets give the number of HOGs present in every genome of the indicated set and absent from the others, where specified. Mpol Clade I (n=8) and Mpol Clade II (n=6) are the two *M. polaris* ecotypes resolved by genome-wide analyses (Fig. 4).

Beyond the shared *Micromonas* core, the HOGs core to *M. polaris* but absent from temperate *Micromonas* formed a polar-specific gene complement several-fold larger (∼2.8x) than the temperate-specific core (Table 2, Fig. 6a). This polar-specific core included the putative ice-binding protein (IBP; PF11999) previously found to be enriched in other polar microalgae.^34^ *M. polaris* shared more HOGs with *M. commoda* than with *M. pusilla*, consistent with *M. commoda* being its closer relative within the genus. The clade-specific cores distinguishing the two ecotypes were small by comparison (Table 2). Most of these clade-specific cores were unannotated (8 of 20 in Clade II), but those with assignable functions included ankyrin repeat domain proteins (3 in Clade I, 8 in Clade II), a regulator of chromosome condensation (RCC1), a histone-fold domain protein, a chromosome segregation protein (SMC), a glycosyl hydrolase, and a viral outer envelope glycoprotein (BLLF1) in Clade II (Extended Data Tables 3 & 4). The asymmetry between the large *M. polaris*-specific core and the small clade-specific cores indicates that polar specialisation is shared across both ecotypes, while the genomic differences between ecotypes lie predominantly in accessory rather than core gene content.

GO enrichment analysis pinpointed the functional axes along which *M. polaris* differs from temperate *Micromonas*. The *M. polaris* pangenome was enriched in terms relating to one-carbon (C1) and folate metabolism (formyltetrahydrofolate deformylase activity, folic acid metabolism, dihydroneopterin aldolase activity, urea catabolism), endosomal trafficking and the cytoskeleton (WASH complex, guanyl-nucleotide exchange factor activity, microtubule motor activity, actin filament binding), ion transport and binding (potassium and monoatomic ion transport, sodium-proton antiporter activity, zinc ion binding, nickel cation binding), protein processing (protein ufmylation, peptidyl-prolyl isomerase activity, protein binding), and stress response (cyanate metabolism, oxidoreductase activity; Fig. 6b). These four functional axes, C1 metabolism, endocytosis, transmembrane transport, and photo-oxidative stress response, organise the *M. polaris*-specific gene complement and motivated the targeted analyses that follow.

#### Four functional themes organise the polar-specific gene set

Phylogenetic context for each polar-specific HOG (Fig. 7, Extended Data Table 5) allowed us to distinguish genes shared with other chlorophytes (green algae of the division Chlorophyta) from candidates for horizontal gene transfer (HGT) by examining the closest non-chlorophyte homologs of each HOG.

**Figure 7.**
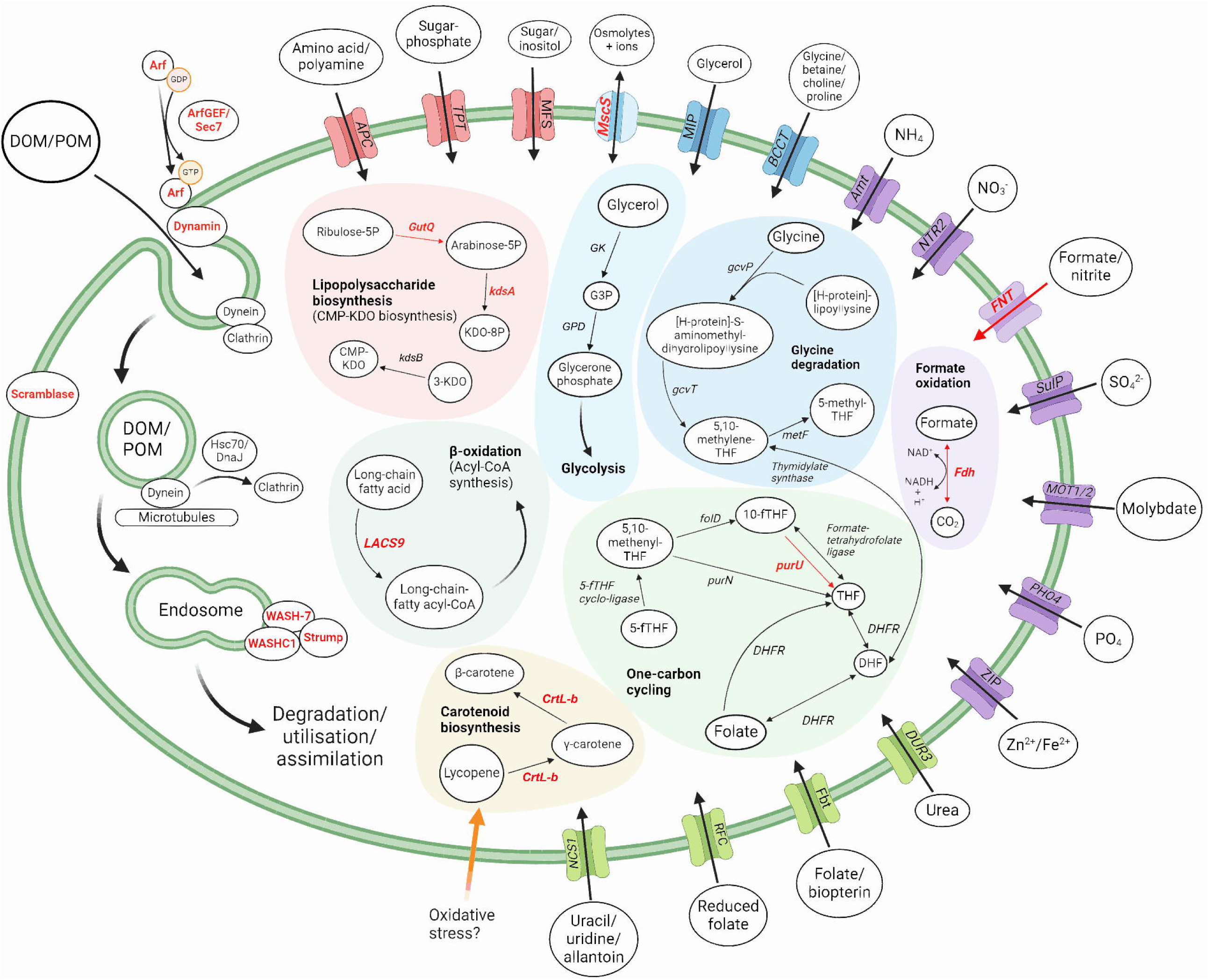
Polar-specific genes integrate into pathways for substrate uptake, alternative carbon metabolism, and osmoregulation. Schematic representation of *M. polaris* cellular pathways involving polar-specific hierarchical orthogroups (HOGs) identified by comparative pangenome analysis (Fig. 6). Gene and protein names in bold red font denote *M. polaris* core HOGs (genes from HOGs shared by all 14 *M. polaris* strains and absent from temperate *Micromonas*); red arrows indicate the corresponding metabolic reactions, which are not encoded in temperate *Micromonas* genomes. Black arrows indicate reactions encoded in both polar and temperate *Micromonas*. Substrates entering or exiting the cell are labelled outside the plasma membrane. **Plasma membrane transporters** (top and right of the cell): an expanded transporter repertoire spans formate/nitrite (FNT), sulfate (SulP), molybdate (MOT1/2), phosphate (PHO4), zinc and iron (ZIP), urea (DUR3), reduced folate (RFC), folate-biopterin (Fbt), nucleobase-cation (NCS1), ammonium (Amt), nitrate (NTR2), amino acid-polyamine-organocation (APC), triose-phosphate (TPT), major facilitator superfamily (MFS), aquaporin (MIP), betaine/carnitine/choline (BCCT), and a mechanosensitive ion channel (MscS). **Bulk fluid-phase uptake of dissolved and fine particulate substrates** (left of the cell): vesicle-mediated internalisation of the surrounding fluid phase, including dissolved organic matter (DOM) and small particulate organic matter (POM) caught up in bulk uptake, involves dynamin, Arf and Arf guanine nucleotide exchange factor (ArfGEF/Sec7), the WASH complex (WASH-7, WASHC1, strumpellin), clathrin, Hsc70/DnaJ chaperones, and phospholipid scramblase, with the long-chain acyl-CoA synthetase LACS9 channelling fatty acids derived from internalised material into β-oxidation. The combination of WASH complex components with other macropinocytosis-associated machinery is consistent with bulk fluid-phase uptake; the absence of the specialised regulators required for selective phagocytosis argues against the regulated ingestion of larger prey particles. **One-carbon (C1) cycling and formate handling**: the polar-specific formyltetrahydrofolate deformylase (purU) releases formate from 10-fTHF, and the polar-specific formate dehydrogenase (Fdh) oxidises imported formate to CO₂ with concomitant NADH generation; the FNT transporter provides the membrane route for formate exchange. The pathway integrates with folate metabolism (DHFR, folD, purN, formate-tetrahydrofolate ligase, 5-fTHF cyclo-ligase) and with glycine degradation (gcvP, gcvT, metF, Thymidylate synthase). **CMP-KDO biosynthesis** (lipopolysaccharide / scale precursor pathway, top left): a near-complete pathway from Ribulose-5P through Arabinose-5P (GutQ, polar-specific), KDO-8P (kdsA, polar-specific), and 3-KDO (kdsB,) to CMP-KDO. **Carotenoid biosynthesis** (bottom left): lycopene β-cyclase (CrtL-b, polar-specific) converts lycopene through γ-carotene to β-carotene, with potential induction under oxidative stress (orange arrow). Glycerol metabolism (GK, GPD) connects to glycolysis. Abbreviations and full descriptions in Extended Data Table 5.

##### C1 metabolism and formate handling

*M. polaris* possessed a near-complete machinery for one-carbon and formate metabolism that temperate *Micromonas* lacked. Three core *M. polaris* genes acted at the centre of this network: a formyltetrahydrofolate deformylase (purU), which converts 10-formyl-THF to THF and releases formate; a formate dehydrogenase (Fdh), which oxidises formate to CO₂ with concomitant reduction of NAD⁺; and a formate/nitrite transporter (FNT), which provides a route for formate exchange across the plasma membrane. *M. polaris* purU shared 57% sequence identity with a homolog in *Coccomyxa subellipsoidea* C-169, another polar chlorophyte microalga.^30^ *M. polaris* Fdh’s closest homolog was in the pennate diatom *Fistulifera solaris*,^50^ a lineage common in polar sea ice. The *M. polaris* FNT had no chlorophyte homologs and instead matched proteins in the pelagophytes *Aureococcus anophagefferens*, *Pelagomonas calceolata,* and the Arctic Pelagophyte *Plocamimonas psychrophila* CCMP2097 (35-36% identity), consistent with acquisition by HGT.

##### Endocytosis and membrane trafficking

Genes for vesicle-mediated nutrient uptake were prominent in the *M. polaris* core, including dynamin, which pinches off vesicles from the plasma membrane,^51^ three subunits of the WASH complex (WASH-7, WASHC1, strumpellin) involved in endosomal cargo sorting,^52^ and a phospholipid scramblase (PLS1). A mechanosensitive ion channel (MscS) lacked any chlorophyte homolog and matched a bacterial single-cell amplified genome from the coast of Maine (USA),^53^ suggesting another likely HGT acquisition.

##### Transmembrane transport beyond formate

*M. polaris* genomes encoded an expanded suite of transporters spanning multiple substrate classes. Beyond the FNT, polar-specific transporters included an amino acid permease, a nucleotide-sugar transporter, a MIP-family aquaporin, and a Pho88 phosphate transporter (Extended Data Table 5). Several of these had non-chlorophyte best hits, raising the possibility that the *M. polaris* transport repertoire was assembled in part through HGT from co-occurring marine taxa.

##### Photo-oxidative stress and the CMP-Kdo pathway

Three core *M. polaris* HOGs implicated in cold and light stress encoded an early light-induced protein (ELIP), a lycopene β-cyclase, and a RuBisCO assembly chaperone (RbcX; Fig. 8). Homologs of all three exist in temperate *Micromonas* but in different HOGs, suggesting that the *M. polaris* copies have diverged sufficiently to cluster separately, possibly reflecting polar-specific functional adaptation. *M. polaris* also possessed a near-complete CMP-Kdo biosynthesis pathway, a feature of the scales present in closely-related Mamiellophyceae *Bathycoccus* spp. and *Dolichomastix* spp.^54^ as well as plant cell walls,^55^ but absent from temperate *Micromonas*. The pathway included a polar-specific arabinose 5-phosphate isomerase (GutQ) whose closest homolog (68% identity) was a protein from the prosthecate marine bacteria *Hyphomonas adhaerens*.^56^

**Figure 8.**
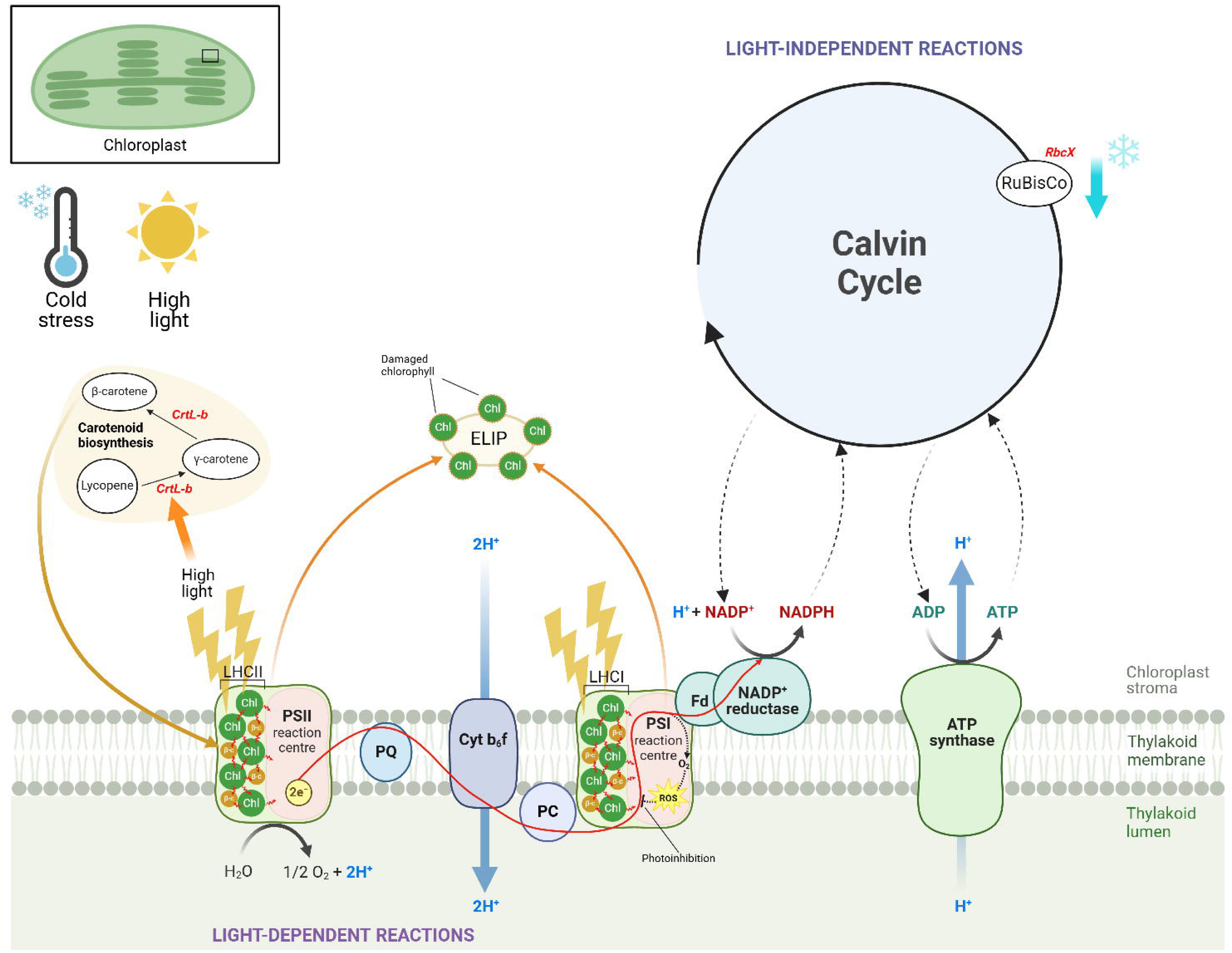
Polar-specific genes for the photo-oxidative stress response integrate into the *M. polaris* chloroplast under combined cold and high-light stress. Schematic representation of the *M. polaris* chloroplast machinery engaged when low temperature and high irradiance occur simultaneously, a combination characteristic of Arctic spring conditions when sea-ice melt exposes acclimated cells to rapidly increasing light. Gene and protein names in bold red font denote *M. polaris* core HOGs (genes from HOGs shared by all 14 *M. polaris* strains and absent from temperate *Micromonas*); these encode three components of the predicted polar-specific photoprotective response. Solid arrows indicate metabolic and electron transport reactions encoded across both polar and temperate *Micromonas*; the orange arrow indicates the proposed transfer of β-carotene toward damaged photosystems under oxidative stress.

Across these four themes, several polar-specific components show phylogenetic signatures consistent with HGT from co-occurring marine taxa, including pelagophytes, methanogens, polar diatoms and marine bacteria.

#### Mpol Clade II is enriched at depth across the Arctic basin

To test whether the depth-partitioning suggested by the strain panel holds at the population level, we quantified the abundance of Mpol Clade I, Mpol Clade II, *M. commoda*, and *M. pusilla* in seven *Tara* Oceans Polar Circle metagenomes^57^ spanning the Northeast Atlantic, Norwegian and Barents Seas, and the Arctic basin (Extended Data Table 6). Each station was sampled at both the surface and the SCM, providing seven paired comparisons of clade abundance across the same water column. Reads were classified against a custom kmer database built from clade-representative genomes (Mpol Clade I: RCC807; Mpol Clade II: RCC2255) and the two temperate *Micromonas* genomes.

Mpol Clade II was the dominant *Micromonas* lineage across all polar-influenced stations, most notably in the Laptev Sea SCM (#189), where 74% of *Micromonas* reads belonged to Clade II and close to 13% of all reads in the sample were classified as *Micromonas* spp. (Fig. 9). Mpol Clade I was consistently present but at lower abundance, with one notable exception: at the Chukchi Sea station (#194), Clade I was relatively more abundant in surface waters than at the SCM. Across the polar stations, the relative abundance of Clade II was higher at the SCM than at the surface in six of seven paired samples, but there was no significant difference in the Clade II to Clade I abundance ratio between SCM and surface samples across the 7 *Tara* stations (Wilcoxon signed-rank test; V=16, *p*=0.81).

**Figure 9.**
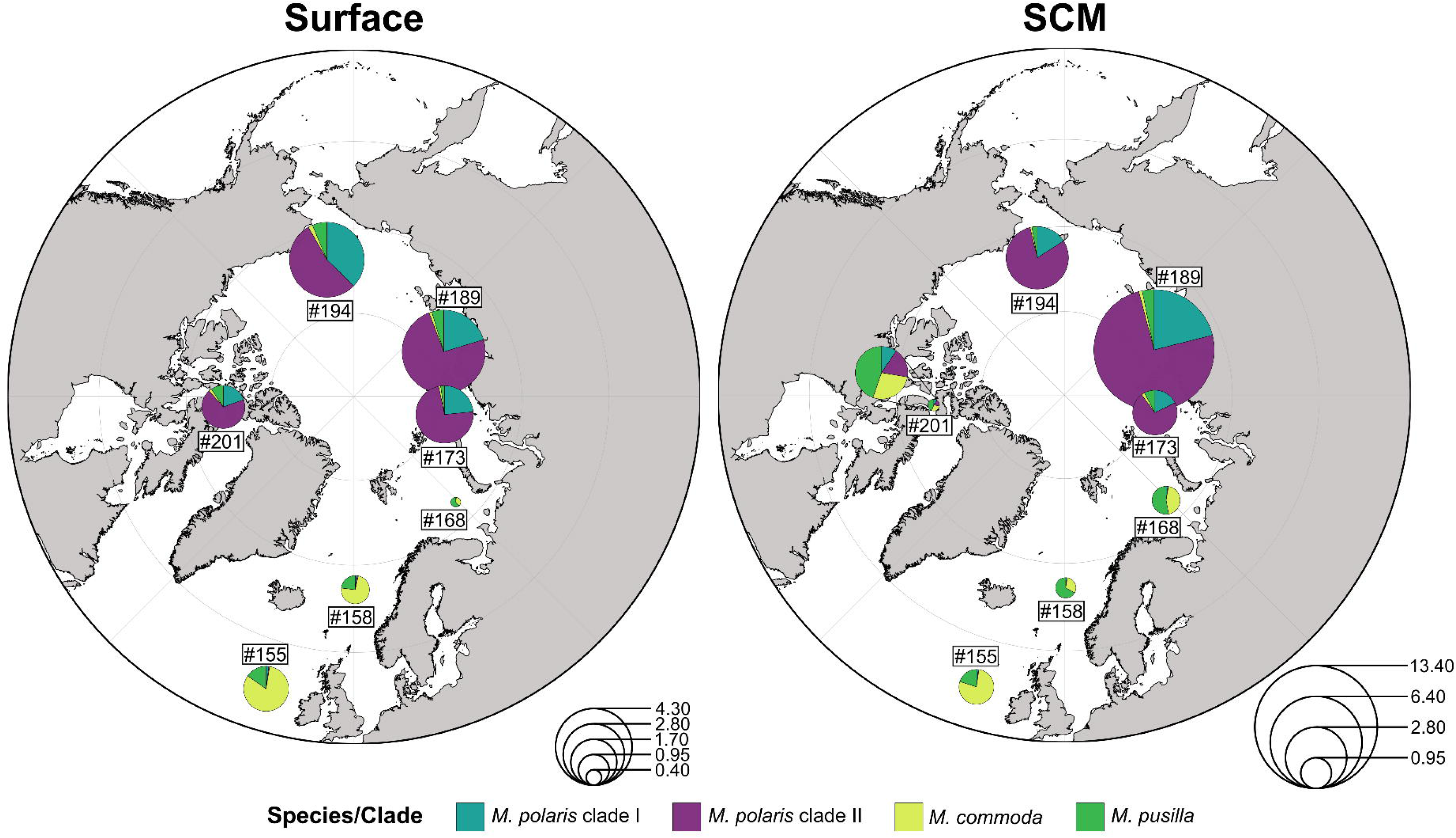
*M. polaris* Clade II dominates Arctic Ocean *Micromonas* communities and is enriched at the subsurface chlorophyll maximum. Biogeographic distribution of *Micromonas* clades and species across seven *Tara* Oceans Polar Circle metagenome samples spanning the Arctic basin, with stations from the Northeast Atlantic, Norwegian Sea, and Barents Sea forming the European Arctic Corridor (stations #155, #158, #168, #173 Kara Sea, #189 Laptev Sea, #194 Chukchi Sea, #201 Canadian Arctic Archipelago). Reads were classified using Kraken2 against a custom database including the 14 *M. polaris* genomes and existing *M. pusilla* and *M. commoda* genomes, with abundance estimates refined by Bracken. Each pie chart shows the *Micromonas* community composition at one station. Pie radius is proportional to the percentage of total metagenomic reads classified as *Micromonas* (scale at lower right of each panel; values are read percentages). Slice colours denote species or clade: dark teal, *M. polaris* Clade I; purple, *M. polaris* Clade II; light green, *M. commoda*; dark green, *M. pusilla*. **a)** Surface samples. *M. polaris* Clade II dominates the *Micromonas* fraction at the central and eastern Arctic stations (#173, #189, #194), with Clade I present at lower abundance. The three European Arctic Corridor stations (#155, #158, #168) instead show temperate *Micromonas* dominance, predominantly *M. commoda*, consistent with the Atlantic-water inflow pattern in this region. **b)** SCM samples. *Micromonas* abundance is generally higher at SCM than at surface (note the larger radius scale relative to panel a), with *M. polaris* Clade II accounting for a higher fraction of the *Micromonas* community at SCM than at surface across the Arctic basin stations.

To test whether clade composition tracked environmental conditions beyond depth categories alone, we related the abundance of each clade in each *Tara* Oceans sample to standardised environmental variables (depth, temperature, salinity, nitrate, and chlorophyll *a*) available for the *Tara* Oceans stations (Extended Data Fig. 8). In exploratory correlation analyses (n=14 samples), Mpol clade I abundance was negatively associated with salinity (Spearman rank correlation; ρ=-0.76, *p*=0.037), while the abundance of Mpol clade II was negatively associated with temperature (Spearman rank correlation; ρ=-0.81, *p*=0.022). These associations are consistent with our observed clade distributions, implying that Mpol clade II may favour the colder, deeper waters below the surface mixed layer, with Mpol clade I preferring fresher, ice-influenced surface waters.

The European Arctic Corridor stations (#155 Northeast Atlantic, #158 Norwegian Sea, #168 Barents Sea) showed distinct signatures. Temperate *Micromonas*, predominantly *M. commoda*, dominated these stations rather than *M. polaris*, and the three clustered together on environmental variables, sharing higher temperatures (5.9 to 11.1 °C), salinities near 35 PSU, and lower chlorophyll *a* than the polar-influenced stations (Extended Data Fig. 8). This biogeographic boundary tracks the documented intrusion of warmer, saltier Atlantic waters into the European Arctic^15,16^ and indicates that temperate *Micromonas* now extend into Atlantic-influenced Arctic margins, where *M. polaris* is correspondingly rare.

To validate *prp8* as a clade marker in environmental sequences, we searched the *Tara* Oceans metatranscriptomes^58^ for *prp8* homologs matching the Mpol Clade I and Clade II consensus sequences (≥99% identity, ≥100 bp alignment). The recovered sequences were placed within Mpol Clade II in a phylogenetic reconstruction (Extended Data Fig. 9) but produced no matches to Mpol Clade I, consistent with the lower environmental abundance of Clade I in the metagenomes. The metagenomic data confirm that the two ecotypes coexist in the Arctic basin and partition vertically with Clade II significantly enriched at depth. The metatranscriptomic data extend this picture to the active community, with Clade II dominating the recovered transcripts and Clade I undetected, consistent with the lower environmental abundance of Clade I in the metagenomes.

## Discussion

*M. polaris* sustains its dominance across the Arctic Ocean as two deeply diverged ecotypes whose distributions are consistently partitioned by depth across the Arctic basin, drawing on a shared polar-specific gene repertoire nearly three times larger than the equivalent set of genes unique to temperate *Micromonas*. The repertoire is enriched in machinery for nutrient acquisition, alternative carbon metabolism, and photo-oxidative stress tolerance, with several components showing phylogenetic signatures consistent with horizontal acquisition from co-occurring marine taxa. Together, these findings reframe how this sentinel species is likely to respond to a warming Arctic.

### Cryptic ecotypes within *M. polaris*

The convergence of *prp8*, genome-wide SNPs, orthogroup-based phylogenies, and *Tara* Oceans metagenomes on the same two-way split supports our interpretation of Mpol Clade I and Mpol Clade II as candidate ecotypes rather than artefacts of any single method, with depth-partitioning predicted by their habitat-of-origin distribution and consistent in environmental data. This pattern places *M. polaris* alongside other marine picoeukaryotes that appear morphologically and phenotypically uniform but contain genetically distinct populations, including the *Bathycoccus prasinos*/*B. calidus* split resolved through genomics^17,18^ and depth-partitioned *Ostreococcus* clades.^19,59^ 18S rRNA, however useful for species delimitation, systematically underestimates ecologically meaningful diversity within marine picoeukaryotes.^22,60,61^ Surveys monitoring *M. polaris* through 18S sequencing therefore collapse two ecologically distinct populations into a single signal and miss differential responses to environmental change. That *prp8* recovers the same two-way split as whole-genome data validates it as a tractable clade marker for environmental work. Together with previous findings in *Bathycoccus*,^45^ this suggests *prp8* may be broadly useful for resolving cryptic diversity within marine microbial eukaryote species, complementing 18S rRNA-based surveys where finer phylogenetic resolution is needed.

### Outlier chromosomes structure mobile and viral element content

Like other Mamiellophyceae, *M. polaris* genomes contain two structurally distinct outlier chromosomes (BOC and SOC; Fig. 5). The conservation of this architecture across all 14 *M. polaris* strains, despite the divergence between Clade I and Clade II, indicates that the two-compartment outlier-chromosome organisation predates the ecotype split and is a stable feature of the species. Within these outlier compartments, TEs and viral insertions concentrate at densities elevated above the genome-wide background, consistent with patterns reported in *Micromonas pusilla* CCMP1545.^37^ The opposite pattern occurs with IEs, which are more common in standard than outlier genomic regions. Clade I genomes carry consistently more introner element families than Clade II (17–20 versus 12–16), the only large-scale architectural difference we observed between the two ecotypes. Whether this difference reflects historical bursts of introner activity in the Clade I lineage, differential selection against introner expansion in Clade II, or contrasting exposure to the viruses and mobile elements that introduce these sequences^37^ cannot be resolved from the data presented here, but the consistency of the difference across all isolates within each clade indicates it is a stable lineage-level feature rather than strain-specific noise. The compartmentalisation of TEs and viral elements into outlier chromosomes also has potential implications for the assembly of the polar-specific gene complement we discuss next: in other Mamiellales, outlier chromosomes are known sites of accelerated gene evolution,^47,62^ and several of the polar-specific HOGs we identified may have entered *M. polaris* genomes through this route.

### A polar-specific genomic toolkit assembled through vertical and lateral inheritance

The 990 polar-specific HOGs shared by both ecotypes encompass functions relevant to flexible carbon and nutrient acquisition under cold, light-limited, highly seasonal conditions, organised around four functional themes: (i) C1 metabolism and formate handling, (ii) endocytosis and membrane trafficking, (iii) expanded transmembrane transport, and (iv) photo-oxidative stress response. The C1 machinery is the most distinctive element of this repertoire, with purU, Fdh, and an FNT transporter together providing a route to import, oxidise, and channel formate into central metabolism. Marine dissolved organic matter contains abundant C1 compounds, and SAR11 and other heterotrophic bacteria use C1 oxidation to generate ATP and reducing power.^63^ Comparable machinery in *M. polaris*, alongside its expanded transporter repertoire and endocytic apparatus, suggests an autotroph capable of substantial heterotrophic supplementation^64^ through bulk fluid-phase uptake (macropinocytosis)^65^ and receptor-mediated endocytosis.^66^

The phylogenetic origin of several polar-specific genes points to HGT from co-occurring marine taxa: an FNT with closest matches in pelagophytes, an Fdh nearest a polar sea ice diatom, a mechanosensitive ion channel matching a coastal bacterial SAG, and a CMP-Kdo pathway component sharing high identity with a marine prosthecate heterotrophic bacterium. Sea ice has been proposed as a hotspot for HGT among algal communities,^35,36^ and recent comparative genomics of Arctic microalgae found that geographically restricted HGT may well be a recurrent feature of Arctic algal genome evolution.^34^ In addition, Arctic algae have been found to possess expanded genomes relative to their temperate counterparts,^34^ a pattern which we observe to a degree in *Micromonas*, where the average *M. polaris* genome size was 22.9 Mbp compared to 21.9 Mbp and 20.9 Mbp in *M. pusilla* CCMP1545 and *M. commoda* RCC299, respectively.^37^ Our gene-by-gene results for *M. polaris* provide a species-level instance of this pattern, identifying specific niche-defining genes whose closest non-chlorophyte homologs lie in independent functional categories of co-occurring marine taxa. Together with growing evidence for HGT in other eukaryotic algae,^34,67,68^ our findings indicate that lateral acquisition from co-occurring marine taxa contributed to the assembly of the *M. polaris* polar niche, alongside vertical inheritance and gene duplication.

The asymmetry between the large polar-specific core (990 HOGs) and the small clade-specific cores (14 in Clade I, 20 in Clade II) indicates that polar specialisation is shared across both ecotypes, with ecotype differentiation likely tuned by accessory and regulatory variation rather than core gene content.

### A genomic basis for non-phagotrophic mixotrophy in *M. polaris*

Whether *M. polaris* combines photosynthesis with heterotrophic nutrition has been debated for over a decade. Phagotrophic mixotrophy was first proposed for the species on the basis of fluorescent bead uptake, with bacterivory invoked as the mechanism allowing growth under low-light conditions.^69^ More recent experimental work across multiple *M. polaris* strains found no evidence of phagocytosis using fluorescent beads or fluorescently labelled bacteria, and proposed that the earlier observations may have reflected surface attachment of prey rather than internalisation.^70^ The two findings remain difficult to reconcile, leaving open the question of how *M. polaris* sustains growth during low-light periods.

Our genomic data clarify the picture: none of the 14 *M. polaris* genomes encode the specialised, regulated machinery required for selective phagocytosis of prey particles, consistent with the experimental absence of bead uptake in recent work.^70^ What the genomes do encode, in the polar-specific gene complement, is an extensive endocytic uptake apparatus: vesicle-mediated internalisation through the WASH complex, dynamin, and phospholipid scramblase PLS1, a combination consistent with non-specific macropinocytosis-style uptake of extracellular fluid in bulk (and dissolved molecules therein) as well as selective, receptor-mediated endocytosis.^65,66,71–73^ This is paired with a transporter repertoire spanning formate, sulfate, molybdate, phosphate, urea, amino acids, folate, and aquaporin-mediated osmolyte exchange. Together, this machinery can internalise dissolved organic matter and any small particulate matter caught through bulk uptake, but lacks the specialised regulators required to recognise, ingest, and digest larger prey particles.

The mixotrophic behaviour previously attributed to *M. polaris* is more parsimoniously interpreted as photosynthesis combined with bulk fluid-phase uptake of dissolved and fine particulate substrates, rather than selective phagotrophy on prey particles. This mechanism is distinct from the regulated phagotrophy of bacterivorous protists such as choanoflagellates or some haptophytes,^74^ which requires machinery the *M. polaris* genomes lack, and may reflect a broader pattern in oligotrophic specialist picoeukaryotes that has been obscured by the historical focus on selective particle ingestion as the diagnostic signature of mixotrophy. For *M. polaris* specifically, the same machinery that supports heterotrophic supplementation in oligotrophic surface waters may also sustain the species through periods when photosynthesis is reduced or absent, providing a genomic basis for survival through the months-long polar night.

### The polar night as a barrier to temperate colonisation

Atlantic-influenced Arctic margins are already dominated by temperate *Micromonas* rather than *M. polaris*. The three *Tara* stations in the European Arctic Corridor (#155 Northeast Atlantic, #158 Norwegian Sea, #168 Barents Sea) clustered together on temperature, salinity, and chlorophyll, with temperate *Micromonas*, predominantly *M. commoda*, dominating the *Micromonas* fraction at all three. These regions correspond to increased Atlantic water inflow over recent decades^75^ and to documented temperate phytoplankton moving north into high latitudes and mixing with Arctic waters.^15,16^ Our results suggest this movement includes the picoplankton fraction.^76,77^ Whether it represents long term replacement or transient summer colonisation depends on how temperate species cope with the Arctic seasonal light regime. Months of continuous darkness during the polar night and prolonged low irradiance during the spring transition will persist regardless of how warm Arctic waters become.

The polar-specific toolkit supports *M. polaris* survival through these light-limited periods, with C1 metabolism, transmembrane transport, and bulk fluid-phase uptake providing alternative carbon and nutrient acquisition routes when photosynthesis is restricted. Temperate *Micromonas* lack this complement, and to date there is no evidence that they could sustain growth through extended darkness.^78^ Temperate *Micromonas* may colonise Arctic margins during the productive light-rich summer, where warming and stratification favour their physiological optima, but retreat or perish during the polar night, leaving *M. polaris* to persist through the dark season. The contrast between summer-suitable and winter-restrictive conditions argues against simple replacement and suggests Arctic margins will become zones of seasonal turnover rather than stable temperate dominance.^78^ Resolving *M. polaris* into ecotypes refines the signal we can read from this species, since the two clades occupy distinct vertical niches and likely differ in their sensitivity to surface warming, stratification, and light penetration.

### Conclusions

Here we found that *M. polaris* consists of two ecotypes, not one, partitioning the Arctic water column by depth and sharing a polar-specific genomic toolkit assembled through a combination of vertical inheritance from chlorophyte ancestors and lateral acquisition from co-occurring marine taxa. Temperate *Micromonas* lack the genetic capacity to survive the polar night, while the polar-specific toolkit allows *M. polaris* to persist through it. Continued Atlantification therefore points to seasonal turnover at Arctic margins rather than longer term replacement. The pangenome and ecotype framework presented here will support work on three open questions: which accessory genes distinguish the ecotypes functionally, how their abundances vary across the polar year, and how ecotype turnover responds as Atlantification reshapes Arctic picoplankton communities.

## Materials & Methods

### Strains and culture conditions

We worked with 14 *M. polaris* strains obtained from the Roscoff Culture Collection (RCC, France; Extended Data Table 1). All cultures were maintained in L1 medium prepared with aged seawater collected from the L4 station in the Western English Channel (50°15.0’N, 4°13.0’W) at 4 °C under continuous low-light conditions for routine maintenance.

### Growth rate and photophysiology experiments

To compare phenotypes across habitats, we selected one representative strain per habitat: RCC2258 (euphotic mixed layer), RCC2309 (subsurface), and RCC5478 (sea ice-influenced). Each strain was grown in triplicate 100 mL cultures under high light (HL; 60 µmol photons m⁻² s⁻¹) and low light (LL; 1.5 µmol photons m⁻² s⁻¹) at 4 °C for 14 days. Light levels were chosen to approximate Arctic summer surface (HL) and SCM (LL) irradiances; LL also approximates light levels beneath thick sea ice.

Starting cell densities (∼1×10⁶ cells mL⁻¹) were standardised across replicates using flow cytometry counts of one-month acclimated cultures. Cultures were sampled daily at the same time (10:00 ± 45 min) for cell counts and photosynthetic quantum yield (*F*_v_/*F*_m_). Cells were counted live within two hours of sampling on a CytoFLEX S flow cytometer (Beckman Coulter, CA, USA), enumerated in 180 µL sample volumes for 3 min at 60 µL/min with the trigger set to red fluorescence. *F*_v_/*F*_m_ was measured from 2 mL subsamples on an AquaPen AP100 fluorometer (Photon Systems Instruments, Czechia). Growth rates were calculated from cell counts on days 6 to 13 using the ‘easy linear’ method in the growthrates v0.8.4 R package.^79^

Effects of strain identity and light treatment on growth rate were tested with Kruskal-Wallis tests; pairwise differences were assessed using Wilcoxon rank-sum tests with Benjamini-Hochberg correction for multiple comparisons.

### Genomic DNA extraction

For each strain, four parallel DNA extractions were performed and pooled to maximise yield. From each two-week-old culture, 10 mL aliquots were centrifuged at 4,000×g for 15 min, and pellets were resuspended in residual medium (∼500 µL) before transfer to 1.5 mL DNA LoBind tubes (Eppendorf, Hamburg, Germany). After a second spin (16,000×g, 3 min), DNA was extracted from each pellet using the DNeasy Plant Mini Kit (Qiagen, Hilden, Germany) following the manufacturer’s instructions, with one modification: a dry centrifugation step (16,000×g, 2 min) was added after the second AW2 wash to remove residual buffer. DNA was eluted in 60 µL of buffer AE preheated to 65 °C.

The four parallel eluates per strain (∼240 µL total) were combined and concentrated using AMPure XP beads (Beckman Coulter, Brea, CA, USA) at a 1.25× ratio, washed twice with 150 µL of 70% ethanol, air-dried for 30 s, and resuspended in 60 µL of nuclease-free water preheated to 55 °C. Final DNA concentrations were quantified using a Qubit4 fluorometer with the 1×dsDNA high-sensitivity assay kit (Invitrogen, Waltham, MA, USA).

### PCR amplification of phylogenetic markers

Two phylogenetic markers were amplified from genomic DNA (Extended Data Table 2): near-full-length 18S rRNA gene fragments using primers targeting conserved regions near the 5’ and 3’ ends,^80^ and a 975 bp region of *prp8* in a conserved exon. *prp8* primers were designed using Primer3Plus^81^ from an alignment of *prp8* genes in the draft RCC807 assembly and the published *M. pusilla* CCMP1545 and *M. commoda* RCC299 genomes (Extended Data Fig. 1), and synthesised by Eurofins Genomics (Cologne, Germany). PCR mixtures and thermocycler programmes are detailed in Extended Data Tables 7 and 8. Amplicons were verified by agarose gel electrophoresis (Extended Data Fig. 10), purified and Sanger-sequenced by Eurofins Genomics, and edited manually in BioEdit v7.7.1.0.^82^ Forward and reverse-complemented reverse reads were merged using pairwise BLASTN alignment.

### Pulsed-field gel electrophoresis (PFGE) karyotyping

Karyotyping of *M. polaris* RCC807 was performed prior to genome assembly to estimate chromosome number. PFGE was run on a 0.8% agarose gel in 0.5× TBE at 6 V/cm with a pulse angle of 120°, using a switching time of 60 s for the first 15 h followed by 120 s for a further 9 h (24 h total run). *Saccharomyces cerevisiae*, *Hansenula wingei*, and concatenated lambda phage served as size standards (Extended Data Fig. 2).

### Genome sequencing

Each *M. polaris* strain was sequenced using a hybrid long-read and short-read approach. Long reads were generated on Oxford Nanopore Technologies (ONT) MinION Mk1C devices using R9.4.1 flow cells, with libraries prepared from 700 to 1,000 ng of pooled genomic DNA using the SQK-LSK109 ligation sequencing kit (ONT, Oxford, UK) and the NEBNext companion module (E7180S; New England Biolabs, Ipswich, MA, USA). Library preparation followed the manufacturer’s protocol, including end-prep, AMPure XP cleanup, adapter ligation, and a final elution in 15 µL of ONT elution buffer. SpotON flow cells (FLO-MIN109) were primed and loaded according to the ONT MinION protocol, and sequencing ran for 72 h.

Short paired-end reads (2×150 bp) were generated on the Illumina NovaSeq platform (San Diego, CA, USA). Library preparation and sequencing were performed by the University of Exeter Sequencing Service using the NEBNext Ultra II FS DNA library prep kit and a NovaSeq SP 300 flow cell.

### Genome assembly

Nanopore signal data were base-called with Guppy v6.5.7+ca6d6af (ONT, UK) using the high-accuracy basecalling model. Read quality was assessed with FastQC v0.12.1.^83^ Long reads were filtered and trimmed with Chopper v0.5.0^84^ (minimum Phred score 8, minimum length 350 bp, first 15 bp trimmed). Short reads were processed with Fastp v0.23.4^85^ using default parameters.

Genomes were assembled from filtered long reads with Flye v2.9.2-b1786^86^ (--nano-raw, --scaffold, --genome-size 20m). Assemblies were polished once with long reads using Medaka v1.9.1^87^ and the r941_min_high_g360 model, then with four iterative rounds of short-read polishing using Pilon v1.24^88^ after mapping with bwa v0.7.17-r1188.^89^ Each polishing round used the previously polished assembly as the new reference. For RCC807, BlobToolKit^90^ was used to filter out non-target contigs (organelles, contaminants) prior to scaffolding by combining sequence length, GC content, coverage, and taxonomic assignment. Scaffolding was performed with LongStitch v1.0.5^91^ for RCC807 and with combinations of LongStitch and ntJoin v1.1.3^92^ for the remaining strains; ntJoin used the polished RCC807 assembly as a scaffolding reference for strains with lower coverage or contiguity (Extended Data Table 9). BUSCO outputs were used to filter contigs prior to scaffolding, retaining only contigs containing universal single-copy ortholog hits. Final assemblies were assessed with QUAST v5.2.0^93^ and BUSCO v5.5.0^94^ in genome mode.

### Genome annotation

Polished nuclear genomes were soft-masked for repeats using RepeatMasker v4.1.2-p1.^95^ For RCC807, existing RNA-seq data for *M. polaris* CCMP2099 (BioProject PRJNA504417) were aligned to the assembly with bwa, and the sorted alignment was provided to BRAKER v3.0.3^96^ (--rnaseq_sets_ids, --rnaseq_sets_dir) to generate a species-specific gene model using AUGUSTUS v3.4.0 and GeneMark-ETP. The resulting RCC807 species model was then used to predict genes in all 14 strains. Predicted proteins were annotated using EggNOG-mapper v2.1.12^97^ for orthology and functional assignment.

IE families were detected using the pipeline of Gozashti *et al*.^98^ IE families were clustered with a sequence similarity threshold of 95% using CD-HIT,^99^ and representative sequences of each cluster were aligned using Mafft. Phylogenetic reconstruction of the IE cluster representatives was carried out with IQ-TREE. The normalised abundance of each IE cluster in each *M. polaris* genome was then plotted as a heatmap, and the *M. polaris* genomes were hierarchically clustered based on their IE abundance profiles. Transposable elements were predicted across whole genomes using EDTA v2.2.2.^100^ Viral-inclusion elements were predicted with ViralRecall v2.^101^ Element densities (IE, TE, viral) were compared between BOC, SOC, and standard chromosomal regions using Wilcoxon rank-sum tests.

### Phylogenetic reconstruction

For 18S rRNA and *prp8* phylogenies, amplicons were aligned using the G-INS-i algorithm in MAFFT v7.258.^102^ 18S amplicons were aligned to an existing reference Mamiellophyceae 18S MSA^103^ using the --addfragments parameter. Alignments were trimmed with TrimAl v1.4.rev15 (-gappyout).^104^ Maximum-likelihood trees were inferred with IQ-TREE v2.2.0.3,^105,106^ with substitution models selected by ModelFinder^107^ and 1,000 non-parametric bootstrap replicates.

For genome-based phylogenies, filtered short reads from each strain were mapped to the RCC807 reference assembly with bwa mem^89^ and sorted with samtools^108^ to produce BAM files. PCR duplicates were removed with MarkDuplicates (Picard).^109^ Variants were called with bcftools^108^ and filtered to retain SNPs only. Variant call format (VCF) files were intersected to retain sites genotyped across all strains, then merged into a single SNP matrix using bcftools query in nucleotide format. The SNP matrix was used as input for IQ-TREE^105,106^ phylogenetic reconstruction with the GTR+ASC model and 1,000 non-parametric bootstrap replicates.

The merged and filtered VCF was also used to generate a BED file with PLINK^110^ for PCA of population structure, also performed in PLINK. Average nucleotide identity between *M. polaris* genomes was computed with FastANI v1.34^111^ using a many-to-many approach, and the resulting matrix was visualised as a heatmap using pheatmap v1.0.12^112^ in R v4.4.1.^113^

### Pangenomic and functional analyses

Non-redundant proteomes from all 14 *M. polaris* strains and the two temperate *Micromonas* genomes (*M. pusilla* CCMP1545, *M. commoda* RCC299) were clustered into hierarchical orthogroups (HOGs) and used to build a species tree with OrthoFinder v3.0,^114^ using pre-computed all-vs-all DIAMOND v2.1.8 blastp searches^115^ (specifying options --ultra-sensitive, --gapopen 8, --gapextend 2, --masking 1 and -e 0.001). Intersections between *Micromonas* HOG sets were visualised with UpSetR^116^ in R v4.4.0.^113^

For Gene Ontology (GO) enrichment analysis, the longest protein from each HOG was extracted as the representative sequence using a custom Python script and annotated with InterProScan v5.70^117^ (--goterms). GO term enrichment in the *M. polaris* pangenome relative to the temperate *Micromonas* pangenome was tested using Fisher’s exact tests on a contingency table of GO term counts, with significance assessed at *p* <0.05. Core *M. polaris* HOGs (present in all *M. polaris* genomes, absent from temperate genomes) were identified, and representative sequences were extracted for metabolic pathway reconstruction using Pathway Tools v23.0.^118^

To assess phylogenetic context of *M. polaris*-specific HOGs and HGT candidates, the longest protein representative of each HOG was retrieved and used as a blastp query against the NCBI non-redundant protein database^119^ (BLOSUM62, e-value threshold 1E-5), and the top 50 hits (according to e-value) were retained for sequence alignment. The same HOG representative sequence was used as a blastp query against the JGI PhycoCosm resource^120^ using the default search parameters (BLOSUM62, e-value threshold 1E-5), and also against the Ocean Gene Atlas v2^58^ (OM-RGCv2+G dastaset) to retrieve environmental homologs (BLOSUM62, e-value threshold 1E-10). The sequences obtained were then aligned using Mafft v7.526 (--auto),^102^ which selected the the L-INS-i algorithm for GutQ, the FFT-NS-i algorithm for MscS and the FFT-NS-2 algorithm for both FNT and Fdh alignments. The resulting alignments were then trimmed using TrimAL v1.5 (-gappyout).^104^ Maximum-likelihood trees were inferred with IQ-TREE v3.1^105,106^ with best substitution models according to BIC selected by ModelFinder^107^ (RAxML model subset; GutQ: LG+R5, MscS: LG+G4, FNT: LG+F+I+R8, Fdh: LG+I+R8) and 1,000 ultrafast bootstrap replicates.^121^

### Biogeographic analysis of *Tara* Oceans metagenomes

To estimate the biogeographic distribution of *Micromonas* species and *M. polaris* clades in the Arctic basin, we analysed metagenomic and metatranscriptomic reads from seven *Tara* Oceans Polar Circle stations^57^ for which both surface and SCM samples were available (Extended Data Table 6). Clade-representative *M. polaris* genomes were selected with Galah v0.4.2^122^ using FastANI v1.34^111^ and a 99% ANI clustering threshold, producing RCC807 (Mpol Clade I) and RCC2255 (Mpol Clade II) as representatives. A custom Kraken2^123,124^ database was built from these two clade representatives plus the *M. pusilla* CCMP1545 and *M. commoda* RCC299 genomes, with the NCBI taxonomy modified using Taxdump Edit^125^ to include the unpublished *M. polaris* representatives. Metagenomic reads were classified against this database with Kraken2, and species- and clade-level abundances were estimated with Bracken v2.9.^126^

Relative abundances were visualised on a polar map with ggOceanMaps v2.2.0^127^ and scatterpie v0.2.4^128^ in R v4.4.0. Standardised environmental data (depth, nitrate, salinity, temperature, chlorophyll *a*) for each *Tara* station and culture isolation site were displayed as a heatmap using ComplexHeatmap v2.20.0.^129^

To validate *prp8* as a marker for *M. polaris* clades in environmental sequences, consensus *prp8* sequences for Mpol Clade I and Mpol Clade II were generated from MSAs of strain amplicons using EMBOSS Cons.^130^ These consensus sequences were used as queries for searches against the *Tara* Oceans Microbiome Reference Gene Catalog v2 metatranscriptomes via the Ocean Gene Atlas v2.0 (https://tara-oceans.mio.osupytheas.fr/ocean-gene-atlas/).^58^ Hits with ≥99% identity over ≥100 bp were retained, added to the strain *prp8* alignment using MAFFT --add with the L-INS-i algorithm, and used for phylogenetic reconstruction with IQ-TREE v2.2.2.7^105,106^ under the RAxML model subset and 100 non-parametric bootstrap replicates.

## Data availability

Raw Illumina and ONT sequencing reads, genome assemblies and genome annotations have been deposited to the ENA under study accession PRJEB129212. Supplementary data are available from Zenodo (doi:10.5281/zenodo.23915246).

## Acknowledgements

We thank Prof Mike Allen for his feedback on an early version of this manuscript. We are grateful to Drs Karen Moore and Michelle Michelsen who shared their expertise in Nanopore sequencing at the beginning of this study, and to Dr Richard Tennant’s lab for allowing us to use their GridION. We also thank Martin Gachenot and the Roscoff Culture Collection team for the algal strains used in this study. This project utilised equipment funded by the Wellcome Trust (Multi-User Equipment Grant award number 218247/Z/19/Z) at the Exeter Sequencing Facility. This research was supported by funding from the Royal Society through a University Research Fellowship (URF\R\180537) and Royal Society Research Fellows Enhancement Award (RGF\R\181012) awarded to Dr Adam Monier, and by the ANR project JCJC PANIC (2022–2026, ANR-21-CE02-0025).

## Author contributions

V.L.N.J.: Conceptualisation, methodology, investigation (lead), formal analysis (lead), visualisation, writing (original draft, review & editing)

K.H.: Formal analysis (supporting), writing (review & editing)

E.D.: Investigation (supporting)

A.C.: Writing (review & editing)

C.L.: Conceptualisation, resources, writing (review & editing)

A.M.: Conceptualisation (lead), methodology, resources, supervision, funding acquisition, writing (original draft, review & editing).

### The dual stress

Cold temperatures slow Calvin cycle activity, particularly RuBisCO carboxylation (downward blue arrow), reducing the demand for NADPH and ATP from the light-dependent reactions. High light continues to drive electron transport through photosystem II (PSII) and photosystem I (PSI), creating an imbalance between excitation rate and downstream electron sink capacity. The resulting over-reduction of the electron transport chain generates reactive oxygen species (ROS) at PSI and causes photoinhibition.

### The polar-specific response

Three *M. polaris* core HOGs encode candidate components of the protective response: (i) the RuBisCO assembly chaperone RbcX is predicted to support RuBisCO folding and activity under cold conditions, partially mitigating the carboxylation bottleneck; (ii) the lycopene β-cyclase CtrL-b acts at two steps of the carotenoid biosynthesis pathway (lycopene → γ-carotene → β-carotene), with β-carotene functioning as an antioxidant and accessory pigment that protects the photosystems from ROS damage; (iii) the early light-induced protein (ELIP) binds chlorophyll molecules released from damaged photosystem subunits, sequestering them during the PSII repair cycle and allowing photosystem reassembly to proceed without further ROS generation from free chlorophylls.

Other components shown are encoded across *Micromonas* species and form the conserved photosynthetic machinery: PSII and PSI reaction centres with their light-harvesting complexes (LHCII, LHCI), the plastoquinone (PQ) and plastocyanin (PC) electron carriers, the cytochrome b₆f complex, ferredoxin (Fd), NADP⁺ reductase, and ATP synthase. The thylakoid membrane partitions the chloroplast stroma (where the Calvin cycle and carotenoid biosynthesis occur) from the thylakoid lumen (where proton accumulation drives ATP synthesis). Abbreviations: Chl, chlorophyll; Chl-c, β-carotene; CtrL-b, lycopene β-cyclase; Cyt b₆f, cytochrome b₆f complex; ELIP, early light-induced protein; Fd, ferredoxin; LHCI/LHCII, light-harvesting complexes I/II; PC, plastocyanin; PQ, plastoquinone; PSI/PSII, photosystem I/II; RbcX, RuBisCO assembly chaperone; ROS, reactive oxygen species.

## References

1. Balzano, S., Marie, D., Gourvil, P. & Vaulot, D. Composition of the summer photosynthetic pico and nanoplankton communities in the Beaufort Sea assessed by T-RFLP and sequences of the 18S rRNA gene from flow cytometry sorted samples. ISME J. 6, 1480–1498 (2012).

2. Berge, J. et al. In the dark: A review of ecosystem processes during the Arctic polar night. Prog. Oceanogr. 139, 258–271 (2015).

3. Li, W. K. W. Annual average abundance of heterotrophic bacteria and *Synechococcus* in surface ocean waters. Limnol. Oceanogr. 43, 1746–1753 (1998).

4. Paulsen, M. L., et al. *Synechococcus* in the Atlantic Gateway to the Arctic Ocean. Front. Mar. Sci. 3, 191 (2016).

5. Lovejoy, C. et al. Distribution, phylogeny, and growth of cold-adapted picoprasinophytes in Arctic seas. J. Phycol. 43, 78–89 (2007).

6. Lovejoy, C., Massana, R. & Pedrós-Alió, C. Diversity and distribution of marine microbial eukaryotes in the Arctic Ocean and adjacent seas. Appl. Environ. Microbiol. 72, 3085–3095 (2006).

7. Monier, A. et al. Oceanographic structure drives the assembly processes of microbial eukaryotic communities. ISME J. 9, 990–1002 (2015).

8. Tragin, M. & Vaulot, D. Novel diversity within marine Mamiellophyceae (Chlorophyta) unveiled by metabarcoding. Sci. Rep. 9, 5190 (2019).

9. Jackson, V. L. N. et al. Vulnerability of Arctic Ocean microbial eukaryotes to sea ice loss. Sci. Rep. 14, 28896 (2024).

10. Bachy, C. et al. Phytoplankton Surveys in the Arctic Fram Strait Demonstrate the Tiny Eukaryotic Alga *Micromonas* and Other Picoprasinophytes Contribute to Deep Sea Export. Microorganisms 10, 961 (2022).

11. Joli, N., Monier, A., Logares, R. & Lovejoy, C. Seasonal patterns in Arctic prasinophytes and inferred ecology of *Bathycoccus* unveiled in an Arctic winter metagenome. ISME J. 11, 1372–1385 (2017).

12. Marquardt, M., Vader, A., Stübner, E. I., Reigstad, M. & Gabrielsen, T. M. Strong Seasonality of Marine Microbial Eukaryotes in a High-Arctic Fjord (Isfjorden, in West Spitsbergen, Norway). Appl. Environ. Microbiol. 82, 1868–1880 (2016).

13. Fadeev, E. et al. Sea ice presence is linked to higher carbon export and vertical microbial connectivity in the Eurasian Arctic Ocean. *Commun*. Biol. 4, 1255 (2021).

14. Priest, T. et al. Atlantic water influx and sea-ice cover drive taxonomic and functional shifts in Arctic marine bacterial communities. ISME J. 17, 1612–1625 (2023).

15. Oziel, L. et al. Faster Atlantic currents drive poleward expansion of temperate phytoplankton in the Arctic Ocean. Nat. Commun. 11, 1705 (2020).

16. Neukermans, G., Oziel, L. & Babin, M. Increased intrusion of warming Atlantic water leads to rapid expansion of temperate phytoplankton in the Arctic. Glob. Change Biol. 24, 2545–2553 (2018).

17. Vannier, T. et al. Survey of the green picoalga *Bathycoccus* genomes in the global ocean. Sci. Rep. 6, 37900 (2016).

18. Bachy, C. et al. Viruses infecting a warm water picoeukaryote shed light on spatial co-occurrence dynamics of marine viruses and their hosts. ISME J. 15, 3129–3147 (2021).

19. Demir-Hilton, E. et al. Global distribution patterns of distinct clades of the photosynthetic picoeukaryote *Ostreococcus*. ISME J. 5, 1095–1107 (2011).

20. Foulon, E. et al. Ecological niche partitioning in the picoplanktonic green alga *Micromonas pusilla*: evidence from environmental surveys using phylogenetic probes. Environ. Microbiol. 10, 2433–2443 (2008).

21. Nolte, V. et al. Contrasting seasonal niche separation between rare and abundant taxa conceals the extent of protist diversity. Mol. Ecol. 19, 2908–2915 (2010).

22. Piganeau, G., Eyre-Walker, A., Grimsley, N. & Moreau, H. How and Why DNA Barcodes Underestimate the Diversity of Microbial Eukaryotes. PLOS ONE 6, e16342 (2011).

23. Clayton, S., Lin, Y.-C., Follows, M. J. & Worden, A. Z. Co-existence of distinct *Ostreococcus* ecotypes at an oceanic front. Limnol. Oceanogr. 62, 75–88 (2017).

24. Simon, N. et al. Revision of the Genus *Micromonas* Manton et Parke (Chlorophyta, Mamiellophyceae), of the Type Species *M. pusilla* (Butcher) Manton & Parke and of the Species *M. commoda* van Baren, Bachy and Worden and Description of Two New Species Based on the Genetic and Phenotypic Characterization of Cultured Isolates. Protist 168, 612–635 (2017).

25. Simmons, M. P. et al. Intron Invasions Trace Algal Speciation and Reveal Nearly Identical Arctic and Antarctic *Micromonas* Populations. Mol. Biol. Evol. 32, 2219–2235 (2015).

26. Belevich, T. A., Ilyash, L. V., Milyutina, I. A., Logacheva, M. D. & Troitsky, A. V. Photosynthetic picoeukaryotes diversity in the underlying ice waters of the White Sea, Russia. Diversity 12, 93 (2020).

27. Freyria, N. J. et al. Salinity tolerance mechanisms of an Arctic Pelagophyte using comparative transcriptomic and gene expression analysis. *Commun*. Biol. 5, 500 (2022).

28. Freyria, N. J. et al. Stress responses in an Arctic microalga (Pelagophyceae) following sudden salinity change revealed by gene expression analysis. *Commun*. Biol. 7, 1084 (2024).

29. Ardyna, M. & Arrigo, K. R. Phytoplankton dynamics in a changing Arctic Ocean. *Nat*. Clim. Change 10, 892–903 (2020).

30. Blanc, G. et al. The genome of the polar eukaryotic microalga *Coccomyxa subellipsoidea* reveals traits of cold adaptation. Genome Biol. 13, R39 (2012).

31. Mock, T. et al. Evolutionary genomics of the cold-adapted diatom *Fragilariopsis cylindrus*. Nature 541, 536–540 (2017).

32. Stephens, T. G. et al. Genomes of the dinoflagellate *Polarella glacialis* encode tandemly repeated single-exon genes with adaptive functions. BMC Biol. 18, 56 (2020).

33. Zhang, Z. et al. Adaptation to Extreme Antarctic Environments Revealed by the Genome of a Sea Ice Green Alga. Curr. Biol. 30, 3330–3341.e7 (2020).

34. Dorrell, R. G. et al. Convergent evolution and horizontal gene transfer in Arctic Ocean microalgae. Life Sci. Alliance 6, e202201833 (2023).

35. Collins, R. E. & Deming, J. W. Abundant dissolved genetic material in Arctic sea ice Part I: Extracellular DNA. Polar Biol. 34, 1819–1830 (2011).

36. Raymond, J. A. & Kim, H. J. Possible Role of Horizontal Gene Transfer in the Colonization of Sea Ice by Algae. PLoS ONE 7, e35968 (2012).

37. Worden, A. Z. et al. Green Evolution and Dynamic Adaptations Revealed by Genomes of the Marine Picoeukaryotes *Micromonas*. Science 324, 268–272 (2009).

38. van Baren, M. J. et al. Evidence-based green algal genomics reveals marine diversity and ancestral characteristics of land plants. BMC Genomics 17, 267 (2016).

39. Massicotte, P. et al. The MALINA oceanographic expedition: how do changes in ice cover, permafrost and UV radiation impact biodiversity and biogeochemical fluxes in the Arctic Ocean? *Earth Syst*. Sci. Data 13, 1561–1592 (2021).

40. Egge, E. et al. An 18S V4 rRNA metabarcoding dataset of protist diversity in the Atlantic inflow to the Arctic Ocean, through the year and down to 1000m depth. *Earth Syst*. Sci. Data 13, 4913–4928 (2021).

41. Randelhoff, A. et al. Seasonality of the Physical and Biogeochemical Hydrography in the Inflow to the Arctic Ocean Through Fram Strait. Front. Mar. Sci. 5, 224 (2018).

42. Bruyant, F. et al. The Green Edge cruise: investigating the marginal ice zone processes during late spring and early summer to understand the fate of the Arctic phytoplankton bloom. *Earth Syst*. Sci. Data 14, 4607–4642 (2022).

43. Massicotte, P. et al. Green Edge ice camp campaigns: understanding the processes controlling the under-ice Arctic phytoplankton spring bloom. *Earth Syst*. Sci. Data 12, 151–176 (2020).

44. Matthes, L. C. et al. Spatial Heterogeneity as a Key Variable Influencing Spring-Summer Progression in UVR and PAR Transmission Through Arctic Sea Ice. Front. Mar. Sci. 7, 183 (2020).

45. Monier, A., Sudek, S., Fast, N. M. & Worden, A. Z. Gene invasion in distant eukaryotic lineages: Discovery of mutually exclusive genetic elements reveals marine biodiversity. ISME J. 7, 1764–1774 (2013).

46. Derelle, E. et al. Genome analysis of the smallest free-living eukaryote *Ostreococcus tauri* unveils many unique features. Proc. Natl. Acad. Sci. 103, 11647–11652 (2006).

47. Blanc-Mathieu, R. et al. Population genomics of picophytoplankton unveils novel chromosome hypervariability. Sci. Adv. 3, e1700239 (2017).

48. Dennu, L. et al. Biological and genomic resources for the cosmopolitan phytoplankton *Bathycoccus*: insights into genetic diversity and function of outlier chromosomes. Plant J. 126, e70982 (2026).

49. Valiadi, M. et al. Three-dimensional genome organisation reveals spatially insulated outlier chromosomes in *Ostreococcus tauri*. BMC Genomics (2026).

50. Tanaka, T. et al. Oil Accumulation by the Oleaginous Diatom *Fistulifera solaris* as Revealed by the Genome and Transcriptome. Plant Cell 27, 162–176 (2015).

51. Ferguson, S. M. & De Camilli, P. Dynamin, a membrane-remodelling GTPase. Nat. Rev. Mol. Cell Biol. 13, 75–88 (2012).

52. Fokin, A. I. & Gautreau, A. M. Assembly and Activity of the WASH Molecular Machine: Distinctive Features at the Crossroads of the Actin and Microtubule Cytoskeletons. Front. Cell Dev. Biol. 9, 658865 (2021).

53. Munson-McGee, J. H. et al. Decoupling of respiration rates and abundance in marine prokaryoplankton. Nature 612, 764–770 (2022).

54. Becker, B., Becker, D., Kamerling, J. P. & Melkonian, M. 2-Keto-Sugar Acids in Green Flagellates: A Chemical Marker for Prasinophycean Scales. J. Phycol. 27, 498–504 (1991).

55. Bar-Peled, M., Urbanowicz, B. R. & O’Neill, M. A. The Synthesis and Origin of the Pectic Polysaccharide Rhamnogalacturonan II – Insights from Nucleotide Sugar Formation and Diversity. Front. Plant Sci. 3, 92 (2012).

56. Weiner, R. M., Melick, M., O’Neill, K. & Quintero, E. *Hyphomonas adhaerens* sp. nov., *Hyphomonas johnsonii* sp. nov. and *Hyphomonas rosenbergii* sp. nov., marine budding and prosthecate bacteria. Int. J. Syst. Evol. Microbiol. 50, 459–469 (2000).

57. Salazar, G. et al. Gene Expression Changes and Community Turnover Differentially Shape the Global Ocean Metatranscriptome. Cell 179, 1068–1083.e21 (2019).

58. Vernette, C. et al. The Ocean Gene Atlas v2.0: online exploration of the biogeography and phylogeny of plankton genes. Nucleic Acids Res. 50, W516–W526 (2022).

59. Rodríguez, F. et al. Ecotype diversity in the marine picoeukaryote *Ostreococcus* (Chlorophyta, Prasinophyceae). Environ. Microbiol. 7, 853–859 (2005).

60. Šlapeta, J., López-García, P. & Moreira, D. Global dispersal and ancient cryptic species in the smallest marine eukaryotes. Mol. Biol. Evol. 23, 23–29 (2006).

61. Leconte, J. et al. Genome Resolved Biogeography of Mamiellales. Genes 11, 66 (2020).

62. Yau, S. et al. Virus-host coexistence in phytoplankton through the genomic lens. Sci. Adv. 6, eaay2587 (2020).

63. Sun, J. et al. One Carbon Metabolism in SAR11 Pelagic Marine Bacteria. PLOS ONE 6, e23973 (2011).

64. Stegemüller, L., Valverde-Pérez, B., Thygesen, A. & Angelidaki, I. Synergistic effects of heterotrophic and phototrophic metabolism for *Haematococcus lacustris* grown under mixotrophic conditions. J. Appl. Phycol. 36, 3175–3186 (2024).

65. Palm, W. Metabolic functions of macropinocytosis. Philos. Trans. R. Soc. B Biol. Sci. 374, 20180285 (2018).

66. McMahon, H. T. & Boucrot, E. Molecular mechanism and physiological functions of clathrin-mediated endocytosis. Nat. Rev. Mol. Cell Biol. 12, 517–533 (2011).

67. Vancaester, E., Depuydt, T., Osuna-Cruz, C. M. & Vandepoele, K. Comprehensive and Functional Analysis of Horizontal Gene Transfer Events in Diatoms. Mol. Biol. Evol. 37, 3243–3257 (2020).

68. Fan, X. et al. Phytoplankton pangenome reveals extensive prokaryotic horizontal gene transfer of diverse functions. Sci. Adv. 6, eaba0111 (2020).

69. McKie-Krisberg, Z. M. & Sanders, R. W. Phagotrophy by the picoeukaryotic green alga *Micromonas* : implications for Arctic Oceans. ISME J. 8, 1953–1961 (2014).

70. Jimenez, V., Burns, J. A., Le Gall, F., Not, F. & Vaulot, D. No evidence of Phago-mixotropy in *Micromonas polaris* (Mamiellophyceae), the Dominant Picophytoplankton Species in the Arctic. J. Phycol. 57, 435–446 (2021).

71. Buckley, C. M., et al. WASH drives early recycling from macropinosomes and phagosomes to maintain surface phagocytic receptors. Proc. Natl. Acad. Sci. 113, E5906–E5915 (2016).

72. Dal Col, J., et al. Phospholipid scramblase 1: a protein with multiple functions via multiple molecular interactors. Cell Commun. Signal. 20, 78 (2022).

73. Kaksonen, M. & Roux, A. Mechanisms of clathrin-mediated endocytosis. Nat. Rev. Mol. Cell Biol. 19, 313–326 (2018).

74. Mitra, A. et al. Defining Planktonic Protist Functional Groups on Mechanisms for Energy and Nutrient Acquisition: Incorporation of Diverse Mixotrophic Strategies. Protist 167, 106–120 (2016).

75. Polyakov, I. V. et al. Greater role for Atlantic inflows on sea-ice loss in the Eurasian Basin of the Arctic Ocean. Science 356, 285–291 (2017).

76. de Sousa, A. G. G. et al. Diversity and Composition of Pelagic Prokaryotic and Protist Communities in a Thin Arctic Sea-Ice Regime. Microb. Ecol. 78, 388–408 (2019).

77. Metfies, K., Appen, W.-J. von, Kilias, E., Nicolaus, A. & Nöthig, E.-M. Biogeography and photosynthetic biomass of Arctic marine pico-eukaryotes during summer of the record sea ice minimum 2012. PLOS ONE 11, e0148512 (2016).

78. Hop, H. et al. Pelagic Ecosystem Characteristics Across the Atlantic Water Boundary Current From Rijpfjorden, Svalbard, to the Arctic Ocean During Summer (2010–2014). Front. Mar. Sci. 6, 181 (2019).

79. Petzoldt, T. growthrates: Estimate Growth Rates from Experimental Data (2022). 10.32614/CRAN.package.growthrates.

80. Moon-van der Staay, S. Y., De Wachter, R. & Vaulot, D. Oceanic 18S rDNA sequences from picoplankton reveal unsuspected eukaryotic diversity. Nature 409, 607–610 (2001).

81. Untergasser, A. et al. Primer3—new capabilities and interfaces. Nucleic Acids Res. 40, e115 (2012).

82. Hall, T. A. BioEdit: a user-friendly biological sequence alignment editor and analysis program for Windows 95/98/NT. Nucleic Acids Symp. Ser. 41, 95–98 (1999).

83. Andrews, S. FastQC: a quality control tool for high throughput sequence data (2010). http://www.bioinformatics.babraham.ac.uk/projects/fastqc.

84. De Coster, W. & Rademakers, R. NanoPack2: population-scale evaluation of long-read sequencing data. Bioinformatics 39, btad311 (2023).

85. Chen, S., Zhou, Y., Chen, Y. & Gu, J. fastp: an ultra-fast all-in-one FASTQ preprocessor. Bioinformatics 34, i884–i890 (2018).

86. Kolmogorov, M., Yuan, J., Lin, Y. & Pevzner, P. A. Assembly of long, error-prone reads using repeat graphs. Nat. Biotechnol. 37, 540–546 (2019).

87. Oxford Nanopore Technologies. Medaka. Oxford Nanopore Technologies (2023).

88. Walker, B. J. et al. Pilon: An Integrated Tool for Comprehensive Microbial Variant Detection and Genome Assembly Improvement. PLOS ONE 9, e112963 (2014).

89. Li, H. & Durbin, R. Fast and accurate short read alignment with Burrows-Wheeler transform. Bioinforma. Oxf. Engl. 25, 1754–1760 (2009).

90. Challis, R., Richards, E., Rajan, J., Cochrane, G. & Blaxter, M. BlobToolKit – Interactive Quality Assessment of Genome Assemblies. G3 Genes Genomes Genet. 10, 1361–1374 (2020).

91. Coombe, L. et al. LongStitch: high-quality genome assembly correction and scaffolding using long reads. BMC Bioinformatics 22, 534 (2021).

92. Coombe, L., Nikolić, V., Chu, J., Birol, I. & Warren, R. L. ntJoin: Fast and lightweight assembly-guided scaffolding using minimizer graphs. Bioinformatics 36, 3885–3887 (2020).

93. Gurevich, A., Saveliev, V., Vyahhi, N. & Tesler, G. QUAST: quality assessment tool for genome assemblies. Bioinformatics 29, 1072–1075 (2013).

94. Simão, F. A., Waterhouse, R. M., Ioannidis, P., Kriventseva, E. V. & Zdobnov, E. M. BUSCO: assessing genome assembly and annotation completeness with single-copy orthologs. Bioinformatics 31, 3210–3212 (2015).

95. Smit, AFA, Hubley, R, & Green, P. RepeatMasker Open-4.0 (2013). http://www.repeatmasker.org.

96. Gabriel, L. et al. BRAKER3: Fully automated genome annotation using RNA-seq and protein evidence with GeneMark-ETP, AUGUSTUS, and TSEBRA. Genome Res. 34, 769–777 (2024).

97. Cantalapiedra, C. P., Hernández-Plaza, A., Letunic, I., Bork, P. & Huerta-Cepas, J. eggNOG-mapper v2: Functional Annotation, Orthology Assignments, and Domain Prediction at the Metagenomic Scale. Mol. Biol. Evol. 38, 5825–5829 (2021).

98. Gozashti, L., et al. Transposable elements drive intron gain in diverse eukaryotes. Proc. Natl. Acad. Sci. 119, e2209766119 (2022).

99. Li, W. & Godzik, A. Cd-hit: a fast program for clustering and comparing large sets of protein or nucleotide sequences. Bioinformatics 22, 1658–1659 (2006).

100. Ou, S. et al. Benchmarking transposable element annotation methods for creation of a streamlined, comprehensive pipeline. Genome Biol. 20, 275 (2019).

101. Aylward, F. O. & Moniruzzaman, M. ViralRecall—A Flexible Command-Line Tool for the Detection of Giant Virus Signatures in ‘Omic Data. Viruses 13, 150 (2021).

102. Katoh, K., Rozewicki, J. & Yamada, K. D. MAFFT online service: multiple sequence alignment, interactive sequence choice and visualization. Brief. Bioinform. 20, 1160–1166 (2019).

103. Marin, B. & Melkonian, M. Molecular Phylogeny and Classification of the Mamiellophyceae class. nov. (Chlorophyta) based on Sequence Comparisons of the Nuclear- and Plastid-encoded rRNA Operons. Protist 161, 304–336 (2010).

104. Capella-Gutiérrez, S., Silla-Martínez, J. M. & Gabaldón, T. trimAl: a tool for automated alignment trimming in large-scale phylogenetic analyses. Bioinformatics 25, 1972–1973 (2009).

105. Nguyen, L.-T., Schmidt, H. A., von Haeseler, A. & Minh, B. Q. IQ-TREE: A fast and effective stochastic algorithm for estimating maximum-likelihood phylogenies. Mol. Biol. Evol. 32, 268–274 (2015).

106. Minh, B. Q. et al. IQ-TREE 2: New Models and Efficient Methods for Phylogenetic Inference in the Genomic Era. Mol. Biol. Evol. 37, 1530–1534 (2020).

107. Kalyaanamoorthy, S., Minh, B. Q., Wong, T. K. F., von Haeseler, A. & Jermiin, L. S. ModelFinder: Fast model selection for accurate phylogenetic estimates. Nat. Methods 14, 587–589 (2017).

108. Danecek, P. et al. Twelve years of SAMtools and BCFtools. GigaScience 10, giab008 (2021).

109. Picard toolkit. Broad Institute, GitHub repository Broad Institute (2019). https://broadinstitute.github.io/picard.

110. Chang, C. C. et al. Second-generation PLINK: rising to the challenge of larger and richer datasets. GigaScience 4, s13742–015-0047–8 (2015).

111. Jain, C., Rodriguez-R, L. M., Phillippy, A. M., Konstantinidis, K. T. & Aluru, S. High throughput ANI analysis of 90K prokaryotic genomes reveals clear species boundaries. Nat. Commun. 9, 5114 (2018).

112. Kolde, R. pheatmap: Pretty Heatmaps (2019). 10.32614/CRAN.package.pheatmap.

113. R Core Team. R: A Language and Environment for Statistical Computing. R Foundation for Statistical Computing (2024). https://www.r-project.org.

114. Emms, D. M. & Kelly, S. OrthoFinder: phylogenetic orthology inference for comparative genomics. Genome Biol. 20, 238 (2019).

115. Buchfink, B., Reuter, K. & Drost, H.-G. Sensitive protein alignments at tree-of-life scale using DIAMOND. Nat. Methods 18, 366–368 (2021).

116. Conway, J. R., Lex, A. & Gehlenborg, N. UpSetR: an R package for the visualization of intersecting sets and their properties. Bioinformatics 33, 2938–2940 (2017).

117. Jones, P. et al. InterProScan 5: genome-scale protein function classification. Bioinformatics 30, 1236–1240 (2014).

118. Karp, P. D. et al. Pathway Tools version 23.0 update: software for pathway/genome informatics and systems biology. Brief. Bioinform. 22, 109–126 (2021).

119. Sayers, E. W. et al. Database resources of the National Center for Biotechnology Information in 2025. Nucleic Acids Res. 53, D20–D29 (2024).

120. Grigoriev, I. V. et al. PhycoCosm, a comparative algal genomics resource. Nucleic Acids Res. 49, D1004–D1011 (2021).

121. Hoang, D. T., Chernomor, O., von Haeseler, A., Minh, B. Q. & Vinh, L. S. UFBoot2: Improving the Ultrafast Bootstrap Approximation. Mol. Biol. Evol. 35, 518–522 (2018).

122. Aroney, S. T. N., Camargo, A. P., Tyson, G. W. & Woodcroft, B. J. Galah: More scalable dereplication for metagenome assembled genomes (2024). 10.5281/zenodo.10526086.

123. Lu, J. et al. Metagenome analysis using the Kraken software suite. Nat. Protoc. 17, 2815–2839 (2022).

124. Wood, D. E., Lu, J. & Langmead, B. Improved metagenomic analysis with Kraken 2. Genome Biol. 20, 257 (2019).

125. Leonard, G. Taxdump Edit (2020). 10.5281/zenodo.3701276.

126. Lu, J., Breitwieser, F. P., Thielen, P. & Salzberg, S. L. Bracken: estimating species abundance in metagenomics data. PeerJ Comput. Sci. 3, e104 (2017).

127. Mikko Vihtakari. ggOceanMaps: Plot Data on Oceanographic Maps using ggplot2 (2024). 10.5281/zenodo.10513701.

128. Guangchuang Yu. scatterpie: Scatter Pie Plot. Bioinformatics Group @ SMU (2024). 10.32614/CRAN.package.scatterpie.

129. Gu, Z., Eils, R. & Schlesner, M. Complex heatmaps reveal patterns and correlations in multidimensional genomic data. Bioinforma. Oxf. Engl. 32, 2847–2849 (2016).

130. Madeira, F. et al. The EMBL-EBI Job Dispatcher sequence analysis tools framework in 2024. Nucleic Acids Res. 52, W521–W525 (2024).

131. Emms, D. M. & Kelly, S. STAG: Species Tree Inference from All Genes. Preprint at 10.1101/267914 (2018).

