## Extended Data for "The pangenome of the Arctic picoeukaryote *Micromonas polaris* resolves cryptic ecotypes and identifies a polar-specific gene repertoire"

Jackson, V.L.N. *et al*.

### Extended Data

**Extended Data Table 1.** Information on the Micromonas polaris strains used in this study. Strains were obtained from the Roscoff Culture Collection (RCC) and originally isolated during four oceanographic campaigns: the North Water Project,[1] the MALINA expedition,[2] the MicroPolar project,[3,4] and the GreenEdge cruise[5] and Ice Camp.[6] Habitat categories (SUB, subsurface; EML, euphotic mixed layer; ICE, sea ice-influenced) reflect the ecological context of isolation rather than nominal isolation depth alone, and are the categories used in Fig. 1.

| Strain | Region of origin | Habitat | Isolation depth (m) | Isolation date | Cruise/expedition |
|---|---|---|---|---|---|
| RCC807 | Baffin Bay | SUB | 55 | April 1998 | North Water Project |
| RCC2242 | Chukchi Sea | EML | 0 | July 2009 | MALINA |
| RCC2244 | Beaufort Sea | EML | 0 | July 2009 | MALINA |
| RCC2252 | Chukchi Sea | EML | 0 | July 2009 | MALINA |
| RCC2255 | Beaufort Sea | EML | 3 | August 2009 | MALINA |
| RCC2258 | Beaufort Sea | EML | 3 | August 2009 | MALINA |
| RCC2304 | Beaufort Sea | SUB | 70 | August 2009 | MALINA |
| RCC2309 | Beaufort Sea | SUB | 56 | August 2009 | MALINA |
| RCC4298 | Greenland Sea | ICE | 20 | March 2014 | MicroPolar |
| RCC4779 | Greenland Sea | SUB | 500 | May 2014 | MicroPolar |
| RCC5422 | Baffin Bay | ICE | 20 | July 2016 | GreenEdge |
| RCC5423 | Baffin Bay | ICE | 0 | July 2016 | GreenEdge |
| RCC5478 | Baffin Bay | ICE | 0 | June 2016 | GreenEdge |
| RCC5480 | Baffin Bay | ICE | 10 | June 2016 | GreenEdge |

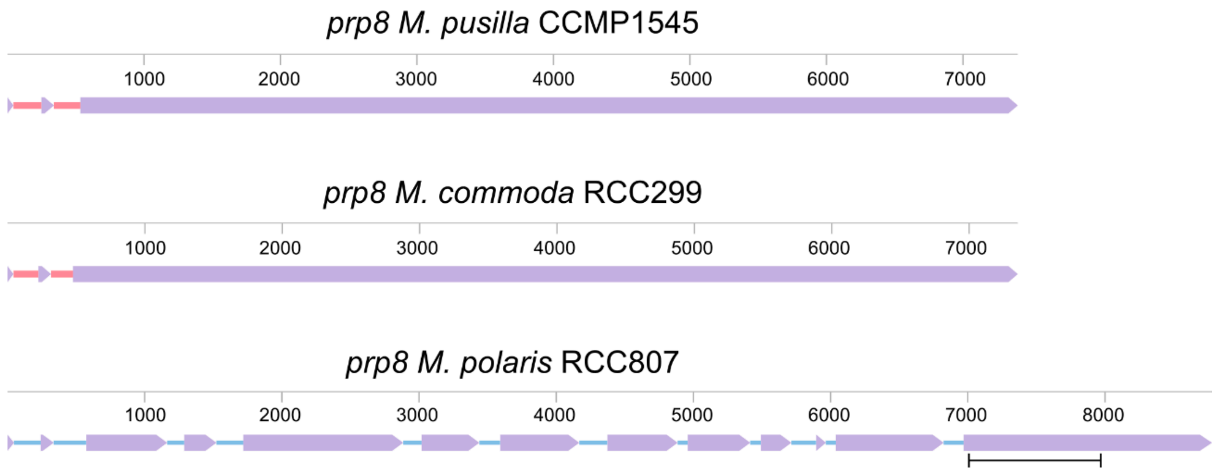


**Extended Data Fig. 1.** *Prp8* gene structure in temperate and polar *Micromonas* spp. Purple segments represent exons, pink segments in CCMP1545 and RCC299 represent introns, and blue segments in RCC807 represent putative introner elements. *M. polaris*-specific primers target a 975 bp sequence in a conserved gene region; the amplicon location on the RCC807 gene is denoted by a black line.

**Extended Data Table 2.** Primer sequences used to generate amplicons of *M. polaris* phylogenetic marker genes. The 18S primers used were previously published by [7], and the *prp8* primers were designed specifically for this study.

| Primer | Sequence |
|---|---|
| 18S-F | 5′-ACCTGGTTGATCCTGCCAG-3′ |
| 18S-R | 5′-TGATCCTTCYGCAGGTTCAC-3′ |
| prp8-8F | 5’-TTCGAGGGTAACCTCACCAC-3’ |
| prp8-8R | 5’-TGCGATCGTGATGAACTTCT-3’ |

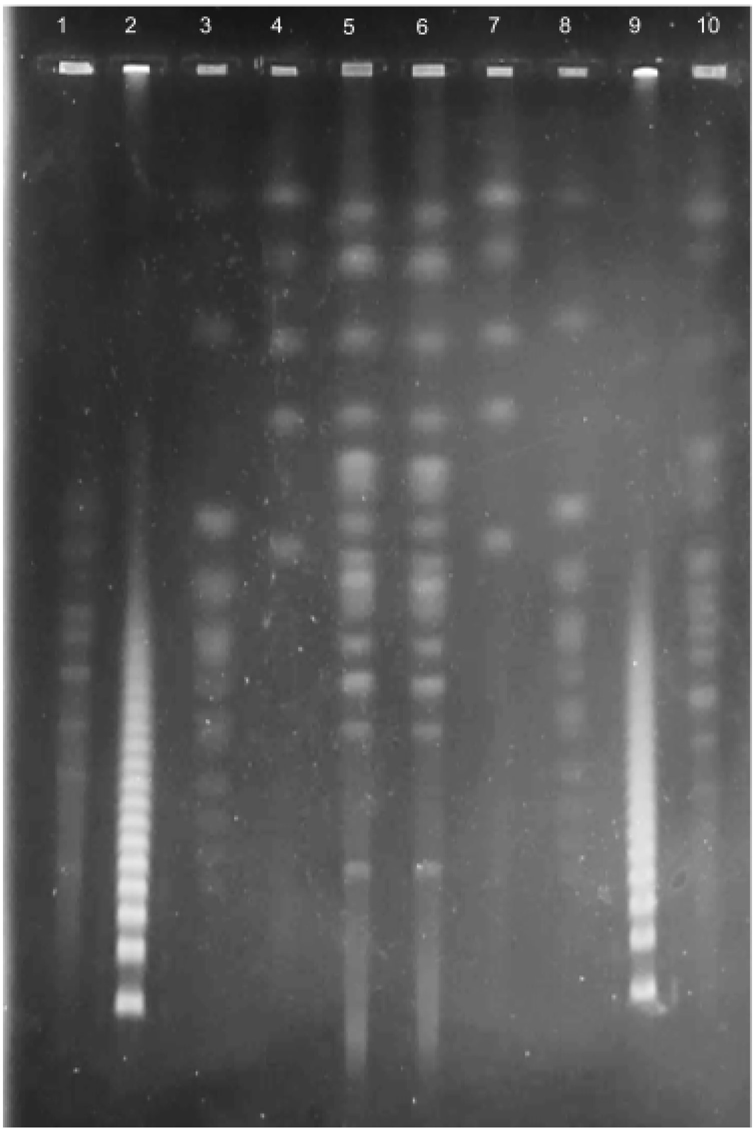


**Extended Data Fig. 2.** PFGE of *M. polaris* RCC807 gDNA. Wells contain: 1, *M. pusilla* CCMP1545; 2 and 9, concatenated lambda phage; 3 and 8, *Saccharomyces cerevisiae*; 4 and 7, *Hansenula wingei*; 5 and 6, *M. polaris* RCC807; 10, *Micromonas* 54-4-A Banyuls.

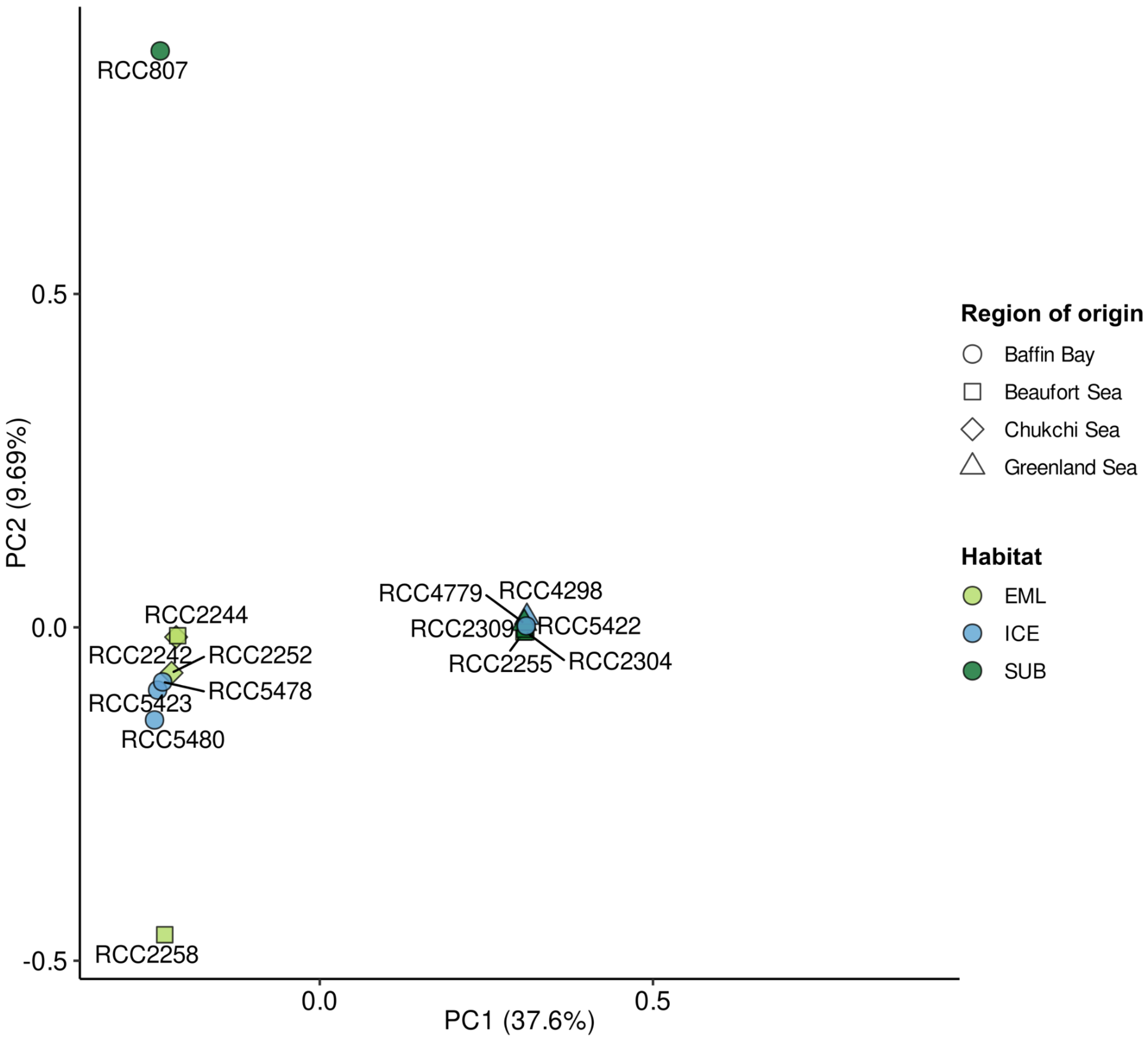


**Extended Data Fig. 3.** PCA plot of *M. polaris* population structure made using PLINK. EML, euphotic mixed layer; ICE, sea ice-associated; SUB, subsurface.

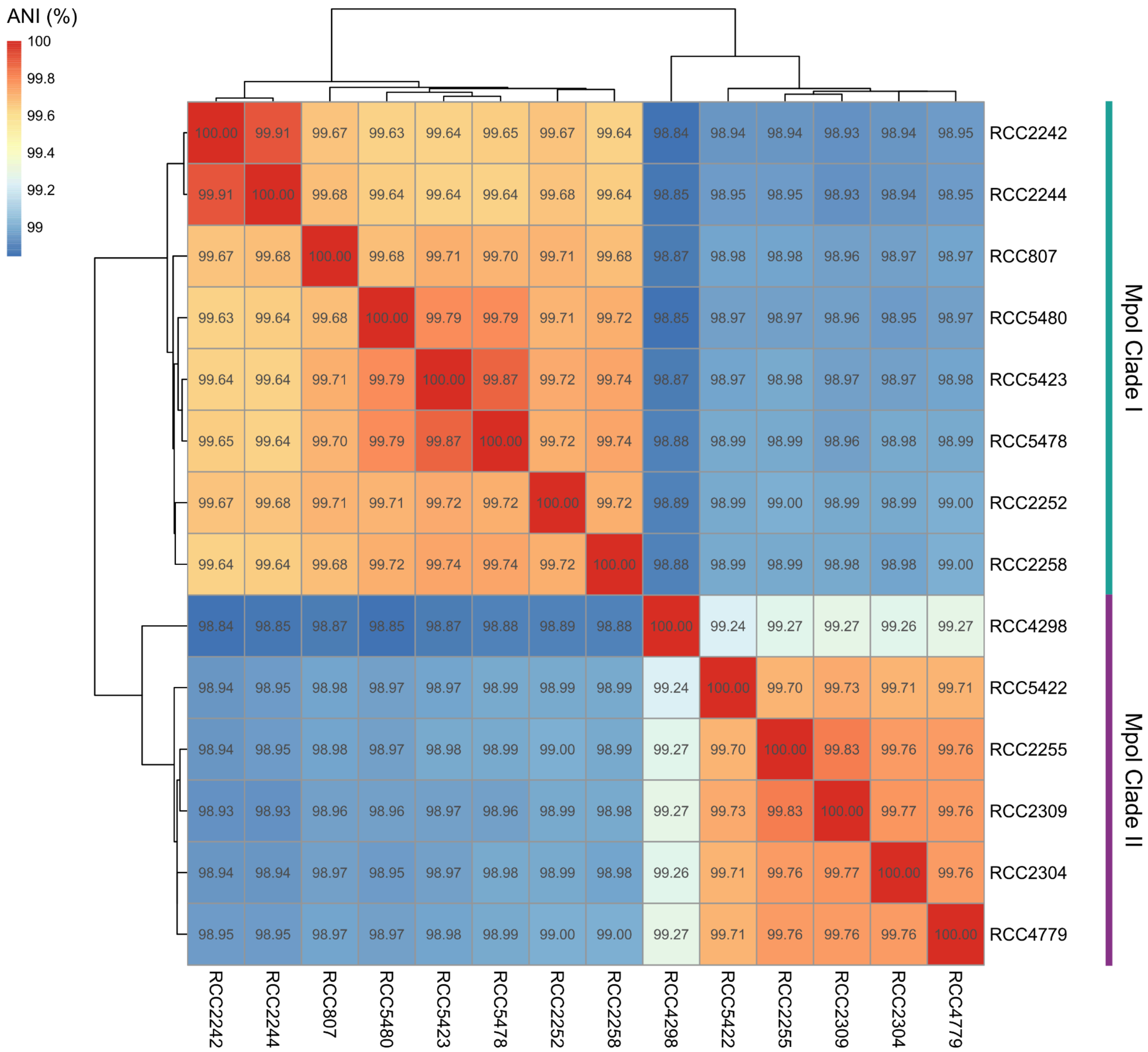


**Extended Data Fig. 4.** ANI heatmap of *M. polaris* genomes. *M. polaris* clades are indicated by coloured bars alongside the heatmap.

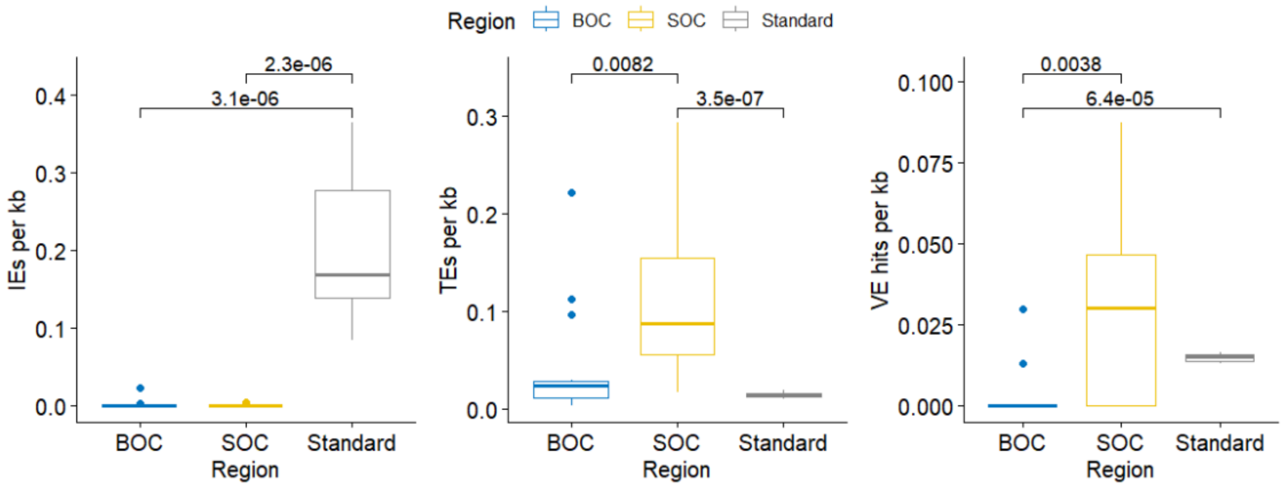


**Extended Data Fig. 5.** Boxplots showing introner element (IE), transposable element (TE), and viral element (VE) abundance per kb of *M. polaris* genomic compartments. BOC, big outlier chromosome; SOC, small outlier chromosome; Standard, standard (non-outlier) genomic region. Significant pairwise comparisons (Wilcoxon) are shown on each plot with their *p* values.

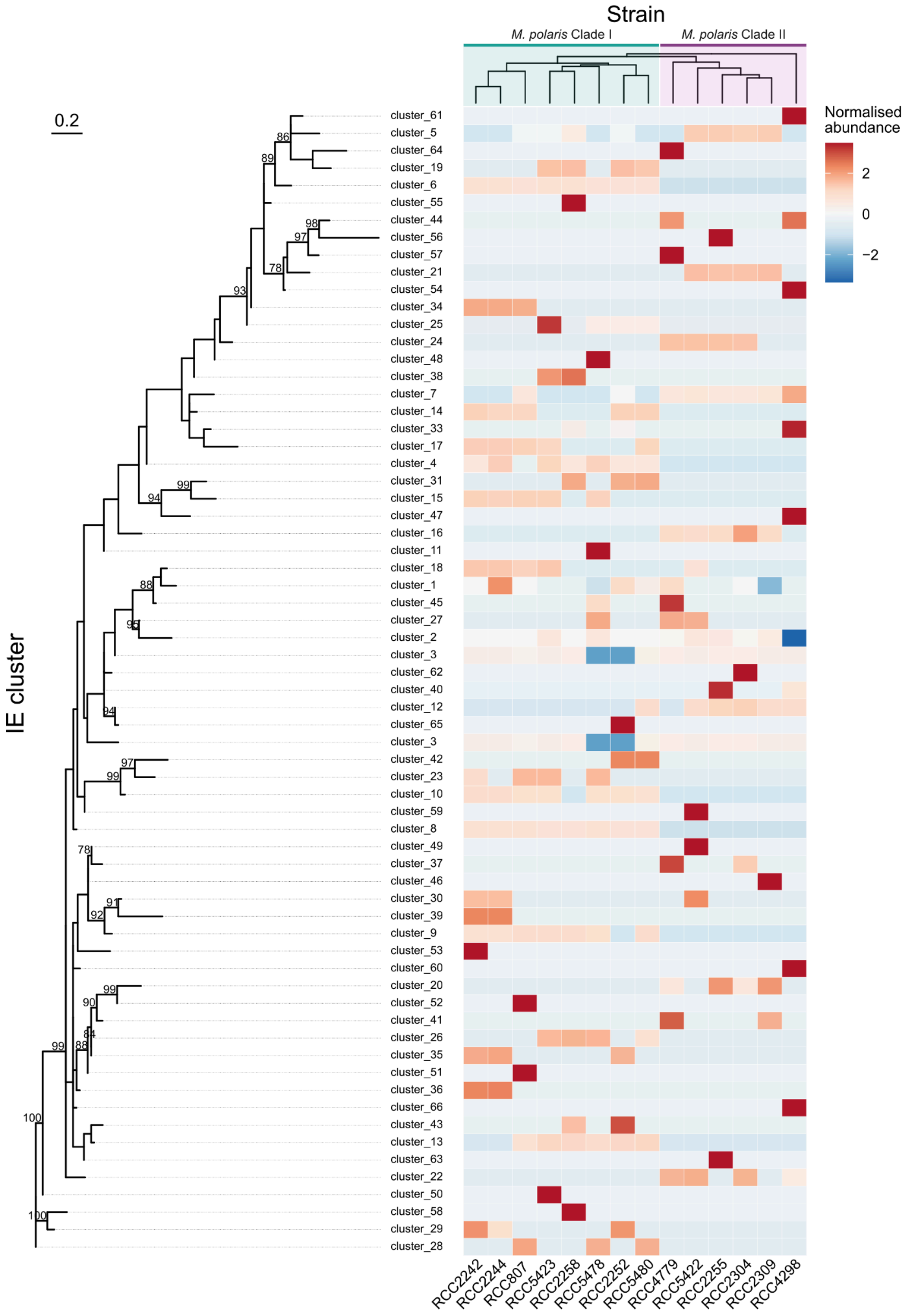


**Extended Data Fig. 6.** Phylogenetic tree and heatmap of IEs in *M. polaris* genomes. IEs across *M. polaris* were clustered with a sequence similarity threshold of 95% using CD-HIT[8] and representative sequences were aligned using Mafft.[9] Phylogenetic reconstruction was carried out with IQ-TREE.[10] Branch support values derived from 1000 ultrafast bootstrap replicates of ≥75% are shown. The heatmap shows the normalised abundance of each IE in each *M. polaris* genome. Columns, representing the genome of each strain, are hierarchically clustered based on the table of normalised IE abundances.

**Extended Data Table 3.** *M. polaris* clade I core HOGs and their annotations.

| HOG | Annotation(s) |
|---|---|
| N0.HOG0009649 | Unknown |
| N0.HOG0009674 | Nar1 domain (iron only dehydrogenase) |
| N0.HOG0009681 | Abhydrolase |
| N0.HOG0009690 | Ankyrin repeats |
| N0.HOG0009728 | Unknown |
| N0.HOG0009738 | MAPK-like serine/threonine protein kinase |
| N0.HOG0009740 | Unknown |
| N0.HOG0009742 | Ankyrin repeats |
| N0.HOG0009743 | Ankyrin repeat |
| N0.HOG0009744 | Unknown |
| N0.HOG0009746 | Unknown |
| N0.HOG0009748 | Unknown |
| N0.HOG0009750 | Unknown |
| N0.HOG0009751 | Tetratricopeptide repeat, beta-lactamase activity |

**Extended Data Table 4.** *M. polaris* clade II core HOGs and their annotations.

| HOG | Annotation(s) |
|---|---|
| N0.HOG0009704 | Ankyrin repeats |
| N0.HOG0009705 | Ankyrin repeats |
| N0.HOG0009730 | Unknown |
| N0.HOG0009731 | Ankyrin repeats |
| N0.HOG0009755 | Unknown |
| N0.HOG0009797 | Histone fold domain found in HF1 & similar proteins |
| N0.HOG0009798 | Glycosyl hydrolase 5 (cellulase A) family |
| N0.HOG0009800 | Ankyrin repeat |
| N0.HOG0009839 | Unknown |
| N0.HOG0009840 | Ankyrin repeats |
| N0.HOG0009841 | Herpes virus major outer envelope glycoprotein (BLLF1) |
| N0.HOG0009842 | Ankyrin repeats |
| N0.HOG0009845 | Unknown |
| N0.HOG0009848 | Unknown |
| N0.HOG0009849 | Ankyrin repeat, chromosome segregation protein SMC |
| N0.HOG0009850 | Unknown |
| N0.HOG0009851 | Regulator of chromosome condensation 1 (RCC1), α-tubulin suppressor (ATS1) domain |
| N0.HOG0009854 | Ankyrin domain |
| N0.HOG0009855 | Unknown |
| N0.HOG0009856 | Unknown |

**Extended Data Table 5.** *M. polaris* core genes putatively involved in $C_1$ metabolism, endocytosis, stress response and degradation/recycling. The hierarchical orthogroup (HOG) of each gene, as well as its representative gene and annotation are shown. Genes are marked as Mpol-specific with an asterisk (□) if there is no homologous gene within the same HOG in temperate *Micromonas*. Phylogenetic background, shown for the Mpol-specific HOGs, shows the likely evolutionary context of each gene based on its best hit(s) in the NCBI non-redundant protein database. Abbreviations: HOG, hierarchical orthogroup; KO, KEGG Orthology code; EC, Enzyme Commission number.

| Mpol-specific | HOG | Description | Pathway(s)/reaction(s) | KO | EC | Phylogenetic background |
|---|---|---|---|---|---|---|
| | | | **$C_1$ cycling & folate metabolism** | | | |
| □ | N0.HOG0009291 | Formyltetrahydrofolate deformylase | | K01433 | 3.5.1.10 | *Coccomyxa viridis* CAK0749969 |
| □ | N0.HOG0002433 | Formate dehydrogenase/ NADH-dependent hydrogenase | Formate oxidation | K00123 K18332 | 1.12.1.3 1.17.1.9 | *Fistulifera solaris* GAX25222 |
| | N0.HOG0007761 | Formate-tetrahydrofolate ligase | | K01938 | 6.3.4.3 | |
| | N0.HOG0004896 | Dihydrofolate reductase/thymidylate synthase | Folate biosynthesis | K13998 | 1.5.1.3 2.1.1.45 | |
| | N0.HOG0000445 | Tetrahydrofolate dehydrogenase/cyclohydrolase | | - | - | |
| | N0.HOG0004108 | 5-formyltetrahydrofolate cycloligase | | K01934 | 6.3.3.2 | |
| | | | **Transmembrane transport** | | | |
| | N0.HOG0002812 | Reduced folate carrier | Vitamin transmembrane transport | K14609, K14610 | - | |
| □ | N0.HOG0009536 | Formate/nitrite transporter | Transmembrane transport of formate/nitrite | K02598 | - | *Plocamimonas psychrophila* CCMP2097 |
| | N0.HOG0001600 | Sulfate permease | Sulfate transport | K18059 | - | |
| □ | N0.HOG0009067 | Amino acid permease | Amino acid transmembrane transport | K03294 | - | Chytrids & allies, bacteria |

| Mpol-specific | HOG | Description | Pathway(s)/reaction(s) | KO | EC | Phylogenetic background |
|---|---|---|---|---|---|---|
| | N0.HOG0006941 | Solute carrier family 35 | Nucleoside-sugar transmembrane transport | K15285 | - | |
| | N0.HOG0006336 | Glycerol-3-phosphate transport | Glycerol-3-phosphate transmembrane transport | K03447 | - | |
| □ | N0.HOG0002051 | Nucleotide-sugar transporter | Nucleotide-sugar transmembrane transport | - | - | *Bathycoccus prasinos*, *Cymbomonas tetramitiformis*, Chlorophyta |
| □ | N0.HOG0009524 | MIP aquaporin family | | K09873 | - | Ochrophyta, Chlorophyta |
| | N0.HOG0000867 | ABC transporter | | K06185 | - | |
| | N0.HOG0003180 | Ammonium transporter | Ammonium uptake | K03320 | - | |
| | N0.HOG0002591 | High-affinity nitrate transporter accessory | High-affinity nitrate uptake | - | - | |
| □ | N0.HOG0007943 | Molybdate transporter MFS superfamily | Molybdenate transport | - | - | *Micromonas commoda*, Chlorophyta |
| □ | N0.HOG0008993 | Phosphate transporter Pho88 | Inorganic phosphate uptake | K14640 | - | Bathycoccaceae, *Aureococcus anophagefferens* |
| | N0.HOG0005292 | Folate-biopterin transporter BT1 | | - | - | |
| | | | **Stress response** | | | |
| □ | N0.HOG0002788 | Early light-induced protein | Photo-oxidative stress response | - | - | Uncultured bacterium, Chlorophyta |
| □ | N0.HOG0009166 | Lycopene β/ε-cyclase | Carotene biosynthesis | - | - | Uncultured bacterium MDC1215515, *Ostreococcus* spp., Phaeophyceae, Chlorophyta |
| | N0.HOG0003976 | Vacuole effluxer ATG22 | Nitrogen starvation response; releases autophagic amino acids | K06902 | - | |
| □ | N0.HOG0008415 | RuBisCO chaperone RbcX | Cold stress response? | - | - | Bathycoccaceae, Streptophyta |
| | N0.HOG0002202 | Mechanosensitive ion channel | Osmotic stress response | K03442 | - | |

| Mpol-specific | HOG | Description | Pathway(s)/reaction(s) | KO | EC | Phylogenetic background |
|---|---|---|---|---|---|---|
| | N0.HOG0006382 | Autophagy protein Apg6 | Heat stress tolerance, autophagy | K08334 | - | |
| | | Ni-containing superoxide dismutase SodN | ROS neutralisation | - | - | |
| **Endocytosis, cytoskeleton and membrane trafficking** | | | | | | |
| □ | N0.HOG0000649 | Dynamin | Vesicle scission | K12476, K12477, K12483 | - | Rhodymeniophycidae, Phaeophyceae, Haptophyta, Dinoflagellata |
| | N0.HOG0008848 | PRA1 | Vesicle-mediated transport | K19496, K20359 | - | |
| | N0.HOG0008563 | Kinesin | Microtubule motor | K08818, K10395, K19626 | 2.7.11.2 2 | |
| | N0.HOG0007630 | Dynein light chain | Microtubule motor | K10411 | - | |
| □ | N0.HOG0008814 | WASH complex subunit 7 | Endosomal WASH complex | K18465 | - | Uncultured bacterium MDA9098370, *Ostreococcus tauri*, Streptophyta |
| | N0.HOG0002042 | WASH complex subunit CCDC53 | | | | |
| □ | N0.HOG0009484 | WASHC1 | Endosomal WASH complex | K18461 | - | *Balaenoptera physalus* KAB0394604 |
| □ | N0.HOG0009352 | Strumpellin | Endosomal WASH complex | K18464 | - | Branchiostoma, Echinodermata |
| □ | N0.HOG0009046 | Long chain acyl-CoA synthetase 9 (LACS9) | | K01897 | 6.2.1.3 | Bathycoccaceae |
| □ | N0.HOG0008023 | Phospholipid scramblase PLS1 | Phospholipid scrambling | - | - | Uncultured bacterium MGY9040087, Chlorophyta, Prymnesiophyceae |
| | N0.HOG0007566 | E3 ubiquitin protein-ligase | | K19045 | 2.3.2.27 | |

| MpoI-specific | HOG | Description | Pathway(s)/reaction(s) | KO | EC | Phylogenetic background |
|---|---|---|---|---|---|---|
| | N0.HOG0006567 | Subunit of the adaptor protein complex 2 (AP-2) | Clathrin-mediated endocytosis | K11824 | - | |
| | N0.HOG0003941 | Subunit of the adaptor protein complex 3 (AP-3) | Vesicle trafficking | K12396 | | |
| | N0.HOG0001997 | ArfGAP | | K12492 | - | |
| □ | N0.HOG0009589 | Sec7 | | K13462 K18442 | - | Temperate *Micromonas* spp., Chlorophyta |
| | N0.HOG0008882 | RasGEF | | K05760, K07520 | - | |
| □ | N0.HOG0009216 | Plexin | | | | Uncultured bacterium MDC1215781, *Ostreococcus tauri* |
| | N0.HOG0003438 | Clathrin heavy chain | Clathrin-mediated endocytosis | K04646 | - | |
| | N0.HOG0006627 | Clathrin light chain | Clathrin-mediated endocytosis | - | - | |
| | N0.HOG0008270 | Hsp70 | Vesicle uncoating | K03283 | - | |
| | N0.HOG0004361 | Syntaxin2 | SNARE | K08488 | | |
| □ | N0.HOG0009420 | Syntaxin | SNARE | K08501 K08503 | | Uncultured bacterium MDA9098086, Streptophyta, Chlorophyta |
| | N0.HOG0005890 | Syntaxin-18 | SNARE | K08492 | | |
| □ | N0.HOG0009204 | COPII-coated ERV/ERGIC | ER-Golgi intermediate compartment protein | K20367 | | *Klebsormidium nitens* GAQ84645, Streptophyta |
| | N0.HOG0003008 | Sorting nexin | | K17925 | - | |
| | | | **Degradation/catabolism** | | | |
| □ | N0.HOG0009027 | 26S proteasome non-ATPase regulatory subunit 10 | Proteasome | K06694 | - | *Ostreococcus tauri* |
| | N0.HOG0002293 | Proteasome subunit Rpn13 | Proteasome | K06691 | - | |

| Mpol-specific | HOG | Description | Pathway(s)/reaction(s) | KO | EC | Phylogenetic background |
|---|---|---|---|---|---|---|
| | N0.HOG0004382 | HslUV peptidase | Proteasome | K01419 | 3.4.25.2 | |
| | N0.HOG0000183 | Proteasome endopeptidase complex | Proteasome | K02738 | 3.4.25.1 | |
| □ | N0.HOG0009397 | CAAX protease | Proteolysis | K07052 | - | Bathycoccaceae, *Coccomyxa* spp. |
| | N0.HOG0001629 | WD40 | Proteasomal ubiquitin-dependent proteolysis | K03362 | - | |
| | N0.HOG0005077 | Patatin/Phospholipase | Triglyceride catabolism | K16816 | 3.1.1.3 | |
| | N0.HOG0004426 | Glycine cleavage system p-protein GCSP | Glycine degradation | K00281 | 1.4.4.2 | |
| | N0.HOG0008524 | Glycine cleavage system t-protein GCST | Glycine degradation | K00605 | 2.1.2.10 | |
| □ | N0.HOG0009049 | GDSL esterase/lipase | Lipid catabolism | - | - | Embryophyta |
| | N0.HOG0008116 | Autophagy protein ATG16 | Autophagy, catabolism | K17890 | - | |
| | N0.HOG0006973 | Autophagy-related protein 3 | | K08343 | | |
| | N0.HOG0002095 | Cysteine protease ATG4 | | K08342 | - | |
| □ | N0.HOG0000705 | Glycerophosphoryl diester phosphodiesterase GDE1 | Glycerophosphodiester degradation | K18696 | 3.1.4.46 | Thalassiosirales |
| □ | N0.HOG0009071 | Ceramidase | Ceramide degradation | K04711 K05848 | 3.5.1.23 | Uncultured bacterium MDB0053579, Myxococcota |
| | N0.HOG0005113 | Autophagy-related protein 9 | | K17907 | - | |
| | N0.HOG0005182 | Autophagy-related protein 101 | | K19730 | - | |
| | N0.HOG0008853 | Urease | Urea degradation | K01427 | 3.5.1.5 | |

**Extended Data Fig. 7.** Phylogenetic reconstructions of 4 horizontal gene transfer (HGT) candidates in *M. polaris*. **a)** Formate dehydrogenase (Fdh); **b)** Formate-nitrite transporter (FNT); **c)** Mechanosensitive ion channel (MscS); **d)** Arabinose 5-phosphate isomerase (GutQ). Tips are coloured according to the phylogenetic origin of sequences (see legend for details) and *M. polaris* sequences are highlighted in yellow on each tree. Scale bars represent substitutions per site and bootstrap support values >75 from 1000 ultrafast bootstrap replicates[11] implemented in IQ-TREE v3.1.3[12] are shown.

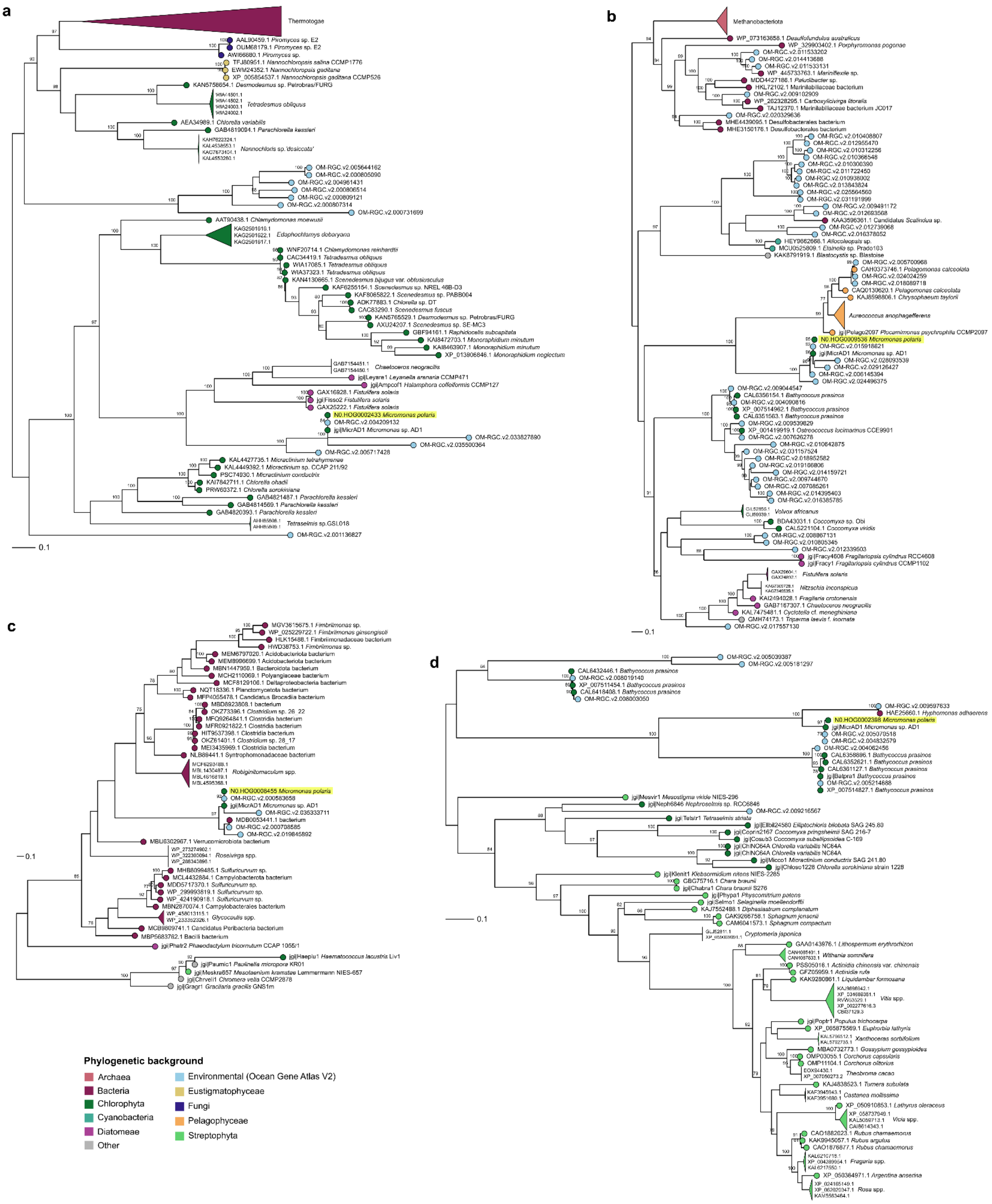

**Extended Data Table 6.** Details of stations from *Tara* Oceans Polar Circle used for biogeographic analysis of Micromonas spp. genotypes.

| ***Tara* station number** | **Region** | **Sampling month** |
|---|---|---|
| 155 | Northeast Atlantic | May |
| 158 | Norwegian Sea | June |
| 168 | Barents Sea | July |
| 173 | Kara Sea | July |
| 189 | Laptev Sea | August |
| 194 | Chukchi Sea | September |
| 201 | Canadian Arctic Archipelago | September |

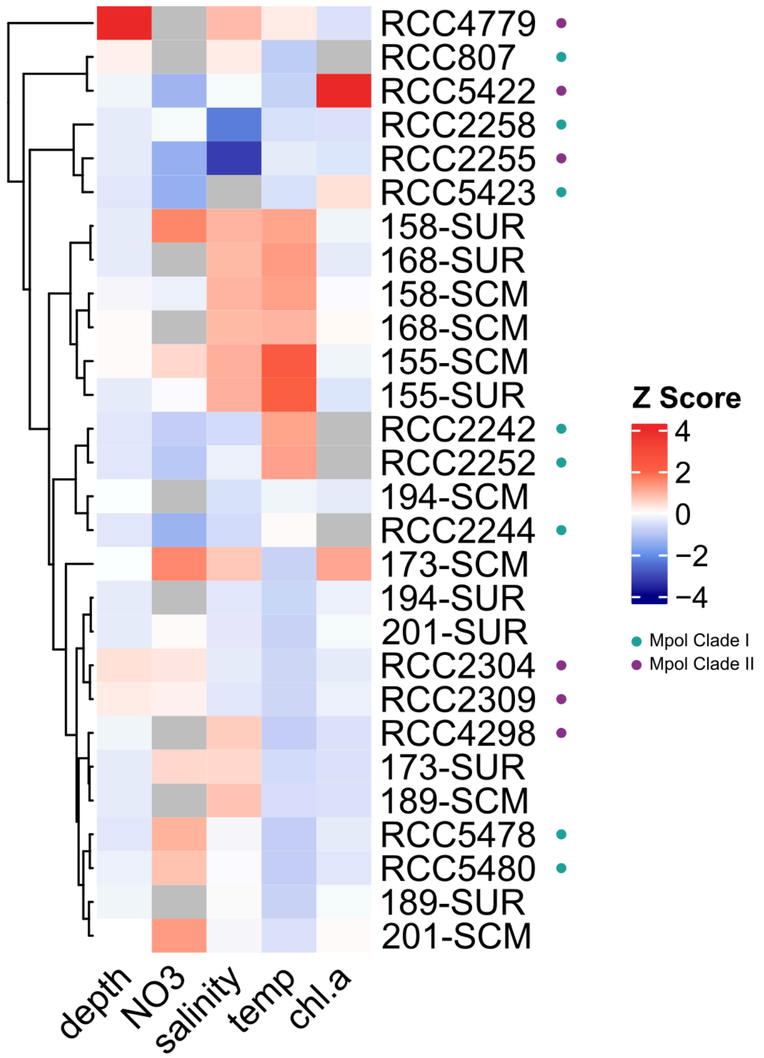


**Extended Data Fig. 8.** Heatmap of standardised (z-score) environmental metadata for each RCC strain isolation site[1–6] and *Tara* station[13] studied here. Coloured circles indicate the clade membership of each *M. polaris* strain.

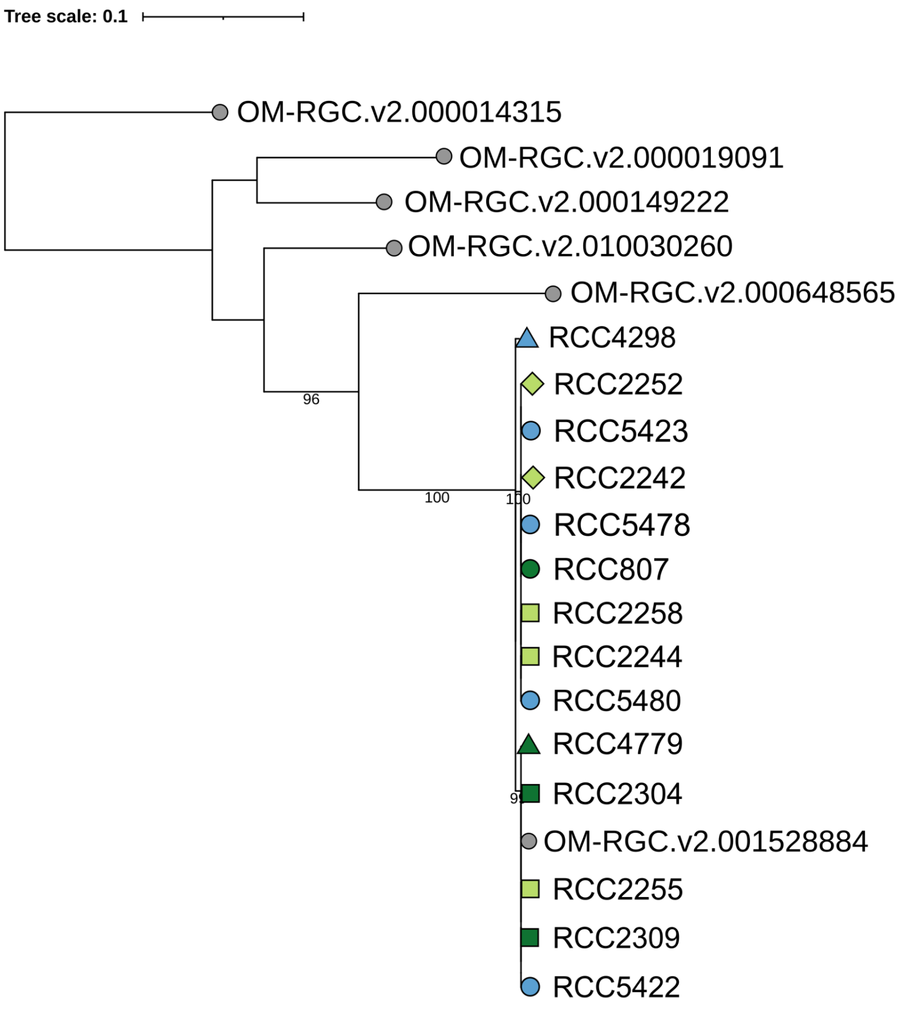


**Extended Data Fig. 9.** Phylogenetic tree of *prp8* amplicons and homolog sequences from the *Tara* Oceans Gene Atlas v2.0 metatranscriptomes.[14]

**Extended Data Table 7.** PCR mixture used to generate amplicons of 18S and *prp8* genes. $dH_2O$ = nuclease-free, deionised water; dNTPs = deoxyribonucleotide mix.

| Component | Volume (µL) |
|---|---|
| $dH_2O$ | 14.9 |
| 5×Reaction buffer | 5 |
| $MgCl_2$ (25 mM) | 1.5 |
| Forward primer (10 µM) | 1 |
| Reverse primer (10 µM) | 1 |
| dNTPs (50 µM) | 0.5 |
| Taq polymerase (5u/µL) | 0.1 |
| Template DNA | 1 |
| **Total reaction volume** | **25** |

**Extended Data Table 8.** Thermocycler programme used for amplification of 18S and *prp8* fragments*.* The denaturation, annealing and elongation steps were repeated 29× for a total of 30 PCR cycles. An annealing temperature of 55 °C was used for 18S primers and 55.8 °C was used for *prp8* primers.

| **Step** | **Temperature (°C)** | **Duration (mm:ss)** |
|---|---|---|
| Initial denaturation | 94 | 03:00 |
| Denaturation | 94 | 00:45 |
| Annealing | 55 (18S) / 55.8 (*prp8)* | 01:00 |
| Elongation | 72 | 02:00 |
| Final elongation | 72 | 05:00 |

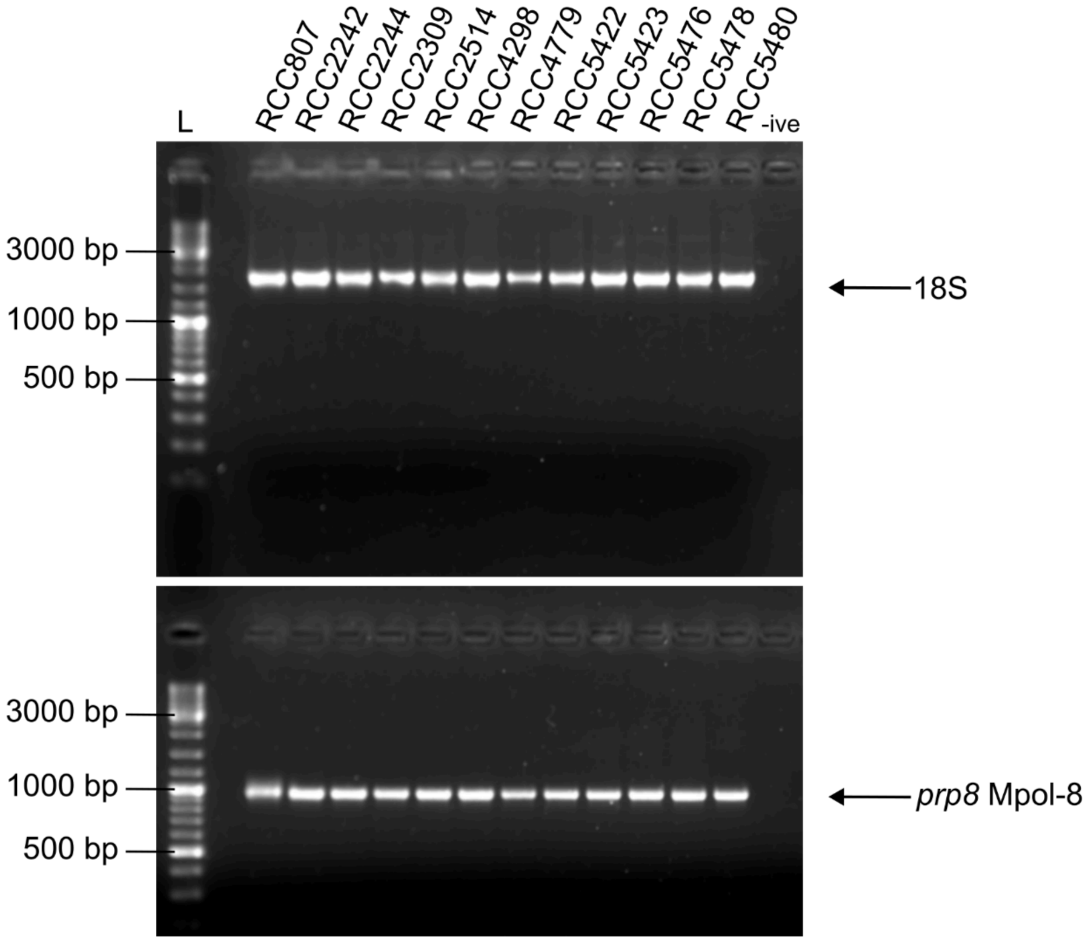


**Extended Data Fig. 10.** Image of 2% agarose gel used to check for successful PCR of 18S and *prp8* amplicons. Upper panel, 18S amplicons; lower panel, *prp8* amplicons; L = ladder; -ive = negative control.

**Extended Data Table 9.** Details of genomic data and assembly statistics for *M. polaris* isolates used in this work. Abbreviations: # ONT reads, number of raw Nanopore reads; # ONT QC reads, number of Nanopore reads after quality control; # Illumina reads, number of raw Illumina reads; # Illumina QC reads, number of Illumina reads after quality control; Ns, percentage of ambiguous bases in genome.

| Strain | # ONT reads | # ONT QC reads | # Illumina reads | # Illumina QC reads | Scaffolding method | Ns (%) |
|---|---|---|---|---|---|---|
| RCC807 | 250353 | 179138 | 43105150 | 42423412 | Longstitch | 0 |
| RCC2242 | 534750 | 520848 | 94931348 | 93519474 | Longstitch×2 | 0.01 |
| RCC2244 | 345219 | 338169 | 64203764 | 63198818 | none | 0 |
| RCC2252 | 159983 | 156141 | 28019210 | 27600892 | Longstitch + ntJoin | 0.10 |
| RCC2255 | 58747 | 57477 | 35862830 | 35331580 | ntJoin + Longstitch×2 + ntJoin | 0.30 |
| RCC2258 | 309704 | 283270 | 23619432 | 23390126 | ntJoin | 0.10 |
| RCC2304 | 361440 | 356473 | 33128968 | 32634732 | Longstitch + ntJoin | 0.22 |
| RCC2309 | 2177701 | 1744109 | 20244664 | 20053378 | ntJoin | 0.30 |
| RCC4298 | 215556 | 195698 | 71351870 | 70034216 | Longstitch + ntJoin | 0.03 |
| RCC4779 | 203629 | 197781 | 88079188 | 87226584 | ntJoin | 0.01 |
| RCC5422 | 279166 | 275485 | 58699028 | 57999676 | none | 0 |
| RCC5423 | 221539 | 218605 | 84895382 | 83668782 | Longstitch + ntJoin | 0.04 |
| RCC5478 | 141376 | 139717 | 50772518 | 50136054 | Longstitch + ntJoin | 0.13 |
| RCC5480 | 333029 | 324837 | 44210492 | 43626180 | Longstitch + ntJoin | 0.26 |

### References


1. Deming, J. W., Fortier, L. & Fukuchi, M. The International North Water Polynya Study (NOW): a brief overview. *Deep Sea Res. Part II Top. Stud. Oceanogr.* **49**, 4887–4892 (2002) doi:10.1016/S0967-0645(02)00168-6.

2. Massicotte, P. *et al.* The MALINA oceanographic expedition: how do changes in ice cover, permafrost and UV radiation impact biodiversity and biogeochemical fluxes in the Arctic Ocean? *Earth Syst. Sci. Data* **13**, 1561–1592 (2021) doi:10.5194/essd-13-1561-2021.

3. Randelhoff, A. *et al.* Seasonality of the Physical and Biogeochemical Hydrography in the Inflow to the Arctic Ocean Through Fram Strait. *Front. Mar. Sci.* **5**, (2018) doi:10.3389/fmars.2018.00224.

4. Egge, E. *et al.* An 18S V4 rRNA metabarcoding dataset of protist diversity in the Atlantic inflow to the Arctic Ocean, through the year and down to 1000m depth. *Earth Syst. Sci. Data* **13**, 4913–4928 (2021) doi:10.5194/essd-13-4913-2021.

5. Bruyant, F. *et al.* The Green Edge cruise: investigating the marginal ice zone processes during late spring and early summer to understand the fate of the Arctic phytoplankton bloom. *Earth Syst. Sci. Data* **14**, 4607–4642 (2022) doi:10.5194/essd-14-4607-2022.

6. Massicotte, P. *et al.* Green Edge ice camp campaigns: understanding the processes controlling the under-ice Arctic phytoplankton spring bloom. *Earth Syst. Sci. Data* **12**, 151–176 (2020) doi:10.5194/essd-12-151-2020.

7. Moon-van der Staay, S. Y., De Wachter, R. & Vaulot, D. Oceanic 18S rDNA sequences from picoplankton reveal unsuspected eukaryotic diversity. *Nature* **409**, (2001) doi:10.1017/CBO9781107415324.004.

8. Li, W. & Godzik, A. Cd-hit: a fast program for clustering and comparing large sets of protein or nucleotide sequences. *Bioinformatics* **22**, 1658–1659 (2006) doi:10.1093/bioinformatics/btl158.

9. Katoh, K., Rozewicki, J. & Yamada, K. D. MAFFT online service: multiple sequence alignment, interactive sequence choice and visualization. *Brief. Bioinform.* **20**, 1160–1166 (2019) doi:10.1093/bib/bbx108.

10. Minh, B. Q. *et al.* IQ-TREE 2: New Models and Efficient Methods for Phylogenetic Inference in the Genomic Era. *Mol. Biol. Evol.* **37**, 1530–1534 (2020) doi:10.1093/molbev/msaa015.

11. Hoang, D. T., Chernomor, O., von Haeseler, A., Minh, B. Q. & Vinh, L. S. UFBoot2: Improving the Ultrafast Bootstrap Approximation. *Mol. Biol. Evol.* **35**, 518–522 (2018) doi:10.1093/molbev/msx281.

12. Nguyen, L.-T., Schmidt, H. A., von Haeseler, A. & Minh, B. Q. IQ-TREE: A fast and effective stochastic algorithm for estimating maximum-likelihood phylogenies. *Mol. Biol. Evol.* **32**, 268–274 (2015) doi:10.1093/molbev/msu300.

13. Guidi, L. *et al.* Environmental context of all samples from the Tara Oceans Expedition (2009-2013), about nutrients in the targeted environmental feature. *In: Tara Oceans Consortium, Coordinators; Tara Oceans Expedition, Participants (2017): Registry of all samples from the Tara Oceans Expedition (2009-2013) [dataset publication series]. PANGAEA, https://doi.org/10.1594/PANGAEA.875582* PANGAEA https://doi.org/10.1594/PANGAEA.875575 (2017) doi:10.1594/PANGAEA.875575.

14. Vernette, C. *et al.* The Ocean Gene Atlas v2.0: online exploration of the biogeography and phylogeny of plankton genes. *Nucleic Acids Res.* **50**, W516–W526 (2022) doi:10.1093/nar/gkac420.
